# A pipeline for facile cloning of antibody Fv domains and their expression, purification, and characterization as recombinant His-tagged IgGs

**DOI:** 10.1101/2024.01.31.578303

**Authors:** Akesh Sinha, Jinha M. Park, Naveed Gulzar, Darpan N. Pandya, Thaddeus J. Wadas, Jamie K. Scott

**Affiliations:** Department of Molecular Biology and Biochemistry, Simon Fraser University, Burnaby, BC V5A 1S6, Canada; Department of Radiology, University of Iowa Hospitals and Clinics, Iowa City, IA 52242, USA; Faculty of Environment, Simon Fraser University, Burnaby BC V5A 1S6 Canada; Faculty of Health Sciences, Simon Fraser University, Burnaby, BC, V5A 1S6, Canada

**Author notes:** Corresponding author: Department of Radiology, 200 Hawkins Drive, 3962 JPP, University of Iowa, Iowa City, IA 52242, USA.

**Keywords:** chimeric antibody, His-tag, affinity purification, single B-cell cloning, surface-plasmon resonance, biolayer interferometry

## Abstract

We report a functional pipeline for facile conversion of variable Fv domains, typically discovered in antibody discovery programs, into chimeric monoclonal antibodies (mAbs). Often, in initial screenings, a set of candidate mAbs is produced in small volumes and purified from supernatant for testing. Our pipeline also simplifies purification of mAbs by using an extended histidine tag (His-10) fused to the C-terminus of the light chain. Both the length of the His-10 and its location have been shown to affect the efficacy of mAb purification using an inexpensive nickel-based resin at neutral pH. Our antibody cloning and purification pipeline, when followed together with detection and affinity measurements, can be smoothly incorporated into an antibody discovery workflow.

## Introduction

Recent technological advancements have increased our ability to discover antigen-specific mAbs at a rapid rate, allowing tremendous progress in the development of novel diagnostics and therapeutics [1, 2]. A common method for discovering new mAbs is the classical hybridoma technology, which requires fusion between immune B cells and a plasmacytoma cell line [3]. The process, however, is inefficient and has low throughput [4, 5]. Other popular approaches to mAb discovery include cloning antibody genes directly from single, antigen-specific B cells [6–11]; synthesizing recombined antibody variable genes deduced from high-throughput sequence data [12–14]; and isolating antigen-binding clones from phage-, yeast- and cell-displayed, naïve [15–18], synthetic [19–23] or immune antibody libraries [24–27]. In addition, antibodies are produced from a variety of species, most typically from mice, rats, rabbits, humans, and more recently, macaques as well as transgenic mice encoding human immunoglobulin germline genes.

As part of our long-term goal of studying B-cell responses against certain HIV glycoproteins, we wanted to test immunogens in different animals and develop a system for cloning DNA encoding the expressed variable heavy and light chain domains (termed VH and VL domains, respectively), together referred as Fv domains, from single, antigen-binding B cells. We wanted to use the same anti-human reagents to detect antibodies bearing these cloned Fv domains; thus, we chose to modify the system of Liao *et al*., which amplifies the recombined V_H_ and V_L_ regions (encoding VH and VL domains, respectively) for cloning and converts them into expression cassettes by overlapping PCR, that encode the constant regions of human heavy (IgG1), kappa or lambda genes [28]. Cells transfected with these constructs have been shown to express mAbs comprising fusions between the cloned Fv domains and human constant regions for both IgG1 heavy and kappa light chains. This allows the use of antibody reagents against human IgGs for detection of any chimeric mAb produced, regardless of the species from which immune B cells are taken. Cloning and transforming Fv domains into full-length, human antibodies also offers the advantages of improved solubility and stability, reduced immunogenicity (in humans) and increased avidity as well as human effector functions.

While Protein A resin is commonly used to purify native or chimeric IgGs [29], its use poses certain limitations: (i) high cost especially when used for semi/high-throughput or industrial scale, (ii) Protein A leaching from the column causing product contamination, (iii) difficulty in sanitizing column material, which may reduce efficiency and potential for reusability, (iv) low pH elution that can reduce activity and/or induce aggregation of the eluted antibody, (v) a limited lifetime compared to other resins [30–35].

Despite advancements in chromatographic procedures for antibody purification, newer non-chromatographic methods are also being explored, owing to the high cost of large-scale production using chromatographic procedures. Some of the non-chromatographic methods for partial or complete antibody purification that has been explored with some success include aqueous two-phase partitioning [36, 37], flocculation and depth filtration [38, 39] and use of co-polymers for antibody precipitation [40]. While these techniques may prove to be economical in large scale antibody purification, their use remains limited due to constraints such as low purity, the need for optimization for different kinds of antibodies and their incompatibility with the rapid small-scale purifications required for screening antibody candidates in high-throughput discovery pipelines.

Peptide tags are better known for their role as antigenic determinants but can also be used for purification of recombinant antibodies. The polyhistidine tag, His-6, is undoubtedly the most widely used affinity tag for purification of recombinant antibody fragments. Immobilized metal affinity chromatography (IMAC) exploits the presence of a His-tag fused to the N- or C-terminus of a recombinant protein, which forms a complex with ionic forms of transition metals like cobalt (Co) or nickel (Ni), allowing the recombinant protein to bind to and then be eluted from a metal column at low pH or with imidazole at neutral pH [41]. His-tag based purification is mostly suitable for recombinant proteins or antibody fragments expressed in *E. coli*, with the purity level of tagged protein reaching more than 80% after IMAC purification [41].

Most full-length antibodies are expressed in insect or mammalian cell cultures that support their folding and post-translation modifications. This presents one of the major limitations in using IMAC for antibody purification, as the higher percentage of histidine residues in insect and mammalian cells can lead to a significant increase in background contaminants that also adhere to the metal column and reduce yield; plus, the presence of non-specific metal-binders in serum can also enhance this effect [41, 42]. As a result, it is not surprising that despite being extensively used, it is uncommon for His-tags to be used for proteins that are expressed in mammalian cell culture. Although, prior studies have reported the use of IMAC for purification of antibodies from complex media including serum [43–45], none of these studies exploited the incorporation of His-tags in the antibody sequence to facilitate antibody purification. Rather, these methods mostly relied on the natural presence of endogenous histidine-containing sequences in antibodies. Some examples include purification of polyclonal goat IgG on Novarose-TREN-Cu^2+^ gel [44], adsorption of human IgG on agarose-IDA- Ni^2+^ resin [45] and purification of humanized murine IgG1 on Tosoh-Haas-TSK-IDA-Ni^2+^ resin [43]. Owing to differences in accessibility of internal histidine residues under various experimental conditions, these purification methods require optimization of column binding and elution conditions for different antibodies and are not generic in nature.

In attempts to overcome these limitations, cell lines having the potential to grow in serum-free media have been developed, allowing one-step IMAC purification of recombinant proteins or antibodies to homogeneity [46, 47]. A modified chelating peptide-tag was successfully used to purify a recombinant antibody from serum-free media by IMAC [48]. However, a major drawback of such systems is the need for unique, adapted cell lines and special media requirements [46]. Advanced media formulations with reduced serum levels and improved cell lines have been developed in recent years which allowing improved yields of recombinant antibodies [49]; but there are no reports of a genetic system that facilitates the use of IMAC to purify recombinant, full-length antibodies even under optimized growth conditions, to the best of our knowledge.

The Strep tag II is another peptide tag that has been employed with some success for one-step purification of recombinant proteins from mammalian cell culture on streptavidin [50–52]; The affinity of Strep tag for streptavidin is weak with K_d_ ∼1 µM [53]. Although the Strep tag generally provides high-purity recombinant proteins, its capacity for binding to its affinity resin is lower [54]. Collectively, these issues indicate a need for alternative antibody purification options that are generic, rapid, economical, reusable, specific, of high affinity and that may be used at different points in antibody discovery pipelines or for large-scale antibody purification.

Here we report a system that simplifies reconstruction of Fv fragments derived from single B cells into human chimeric mAbs and facilitates their reliable expression in mammalian cell culture. Our cloning system also incorporates fusion to an extended His-tag, His-10, located at a site that supports high-yield purification of mAbs from Ni-NTA (Nitrilotriacetic acid) affinity resin directly from cell culture media. As a proof-of-concept of this system, we immunized rabbits with filamentous bacteriophage and created phage-specific, rabbit-human IgG1 mAbs by cloning cDNAs encoding expressed VH and VL domains from single, antigen-specific B cells into our expression system. The culture supernatants from a set of candidate clones were used to purify phage-specific, rabbit-human chimeric mAbs by one-step affinity purification. We also show that purified, recombinant human chimeric mAb bearing His-10 can be immobilized to a Ni-NTA biosensor surface for determination of affinity constant for antigen *via* SPR. While the techniques described here are well established, their unique combination in this application provides a facile system for mAb production, screening, purification, and testing.

## Materials and methods

### Cloning V_H_ and V_L_ sequences to generate human chimeric antibody genes

The segments of antibody genes encoding their VH and VL domains were ligated to human heavy and light chain constant region genes using size overlap extension PCR as shown in Figure 1. Phusion DNA polymerase (Thermo Fisher Scientific, Waltham, MA, USA) was used for size overlap PCR following the manufacturer’s recommendations. Linear cassettes containing a cytomegalovirus promoter (P_CMV_) with leader sequence or human kappa chain with a terminal poly A sequence were obtained as a kind gift from Dr. Hua-Xin Liao [28]. A linear expression cassette encoding IgG1 C_H_ domains and a terminal poly A sequence was derived from a plasmid encoding the heavy chain of mAb 10E8 (kindly donated to the NIH AIDS Reagent Program by Drs. Jinghe Huang, Leo Laub and Mark Connors, NIAID, NIH, Bethesda, Maryland, USA). V_H_ and V_L_ gene segments used in this study were derived either from well-characterized hybridoma cell lines using RT-PCR (for murine mAbs 8H11 and 17/9) or from single B-cell cloning (for mAb 238A1). Reverse transcriptase reactions were performed on mRNAs isolated from hybridoma cell lines using RNeasy Micro kit (Qiagen, Hilden, Germany), followed by Superscript first strand synthesis reaction (Thermo Fisher Scientific, Waltham, MA, USA) with oligo-dT priming, following manufacturer’s recommendations. This was followed by PCR amplification of the antibody V_H_ and V_L_ regions using Phusion DNA polymerase. The amplified V_H_ and V_L_ regions were converted to chimeric antibody heavy and light chain genes using size-overlap extension PCR with flanking linear cassettes encoding P_CMV_ and corresponding heavy- or light-chain constant regions. The primers used are listed in Table 1. The resulting human chimeric heavy and light chain genes were separately cloned in pJET 1.2/blunt vector ( Thermo Fisher Scientific, Waltham, MA, USA). The insertion of DNA encoding His-tag was performed using appropriate primers (Table 1) and a PfuUltra high-fidelity DNA polymerase (Agilent Technologies, Santa Clara, CA, USA) PCR reaction.

**Figure 1.**
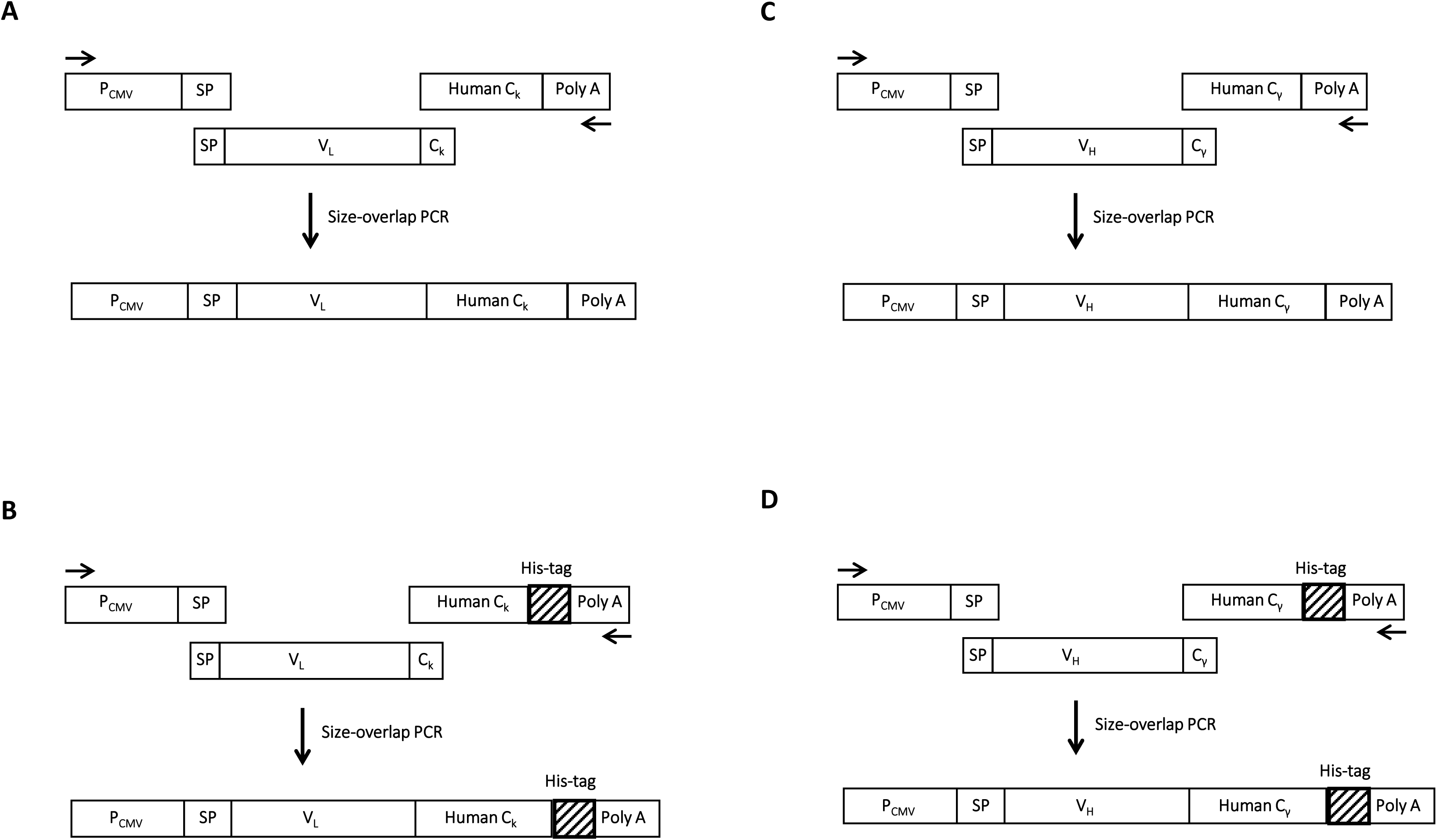
Schematic diagram for generation of human chimeric antibody heavy and light chain genes from overlapping linear expression cassettes: Different panels represent the assembly of full-length genes by size-overlap extension for human chimeric kappa light chain in the absence or presence of C-terminal His-tag (A and B) and for human chimeric gamma 1 heavy chain in the absence or presence of a C-terminal His-tag (C and D). The P expression cassette containing P_CMV_, and the signal peptide (SP) coding sequence is common to all gene assemblies. The V cassette with V_H_ or V_L_ coding sequences are user defined where overlapping sequence of SP and C_H_/Cκ are appended at the 5’ and 3’ ends, respectively for overlap with the other two cassettes. The C cassette encoding the antibody light or heavy chain constant regions also contains a poly A sequence (Poly A). The same forward and reverse primers (Fw C-fragment and Rev H/K fragments, as shown by arrows) were used in size overlap PCR for assembling all chimeric light and heavy chain genes.

**TABLE 1.**
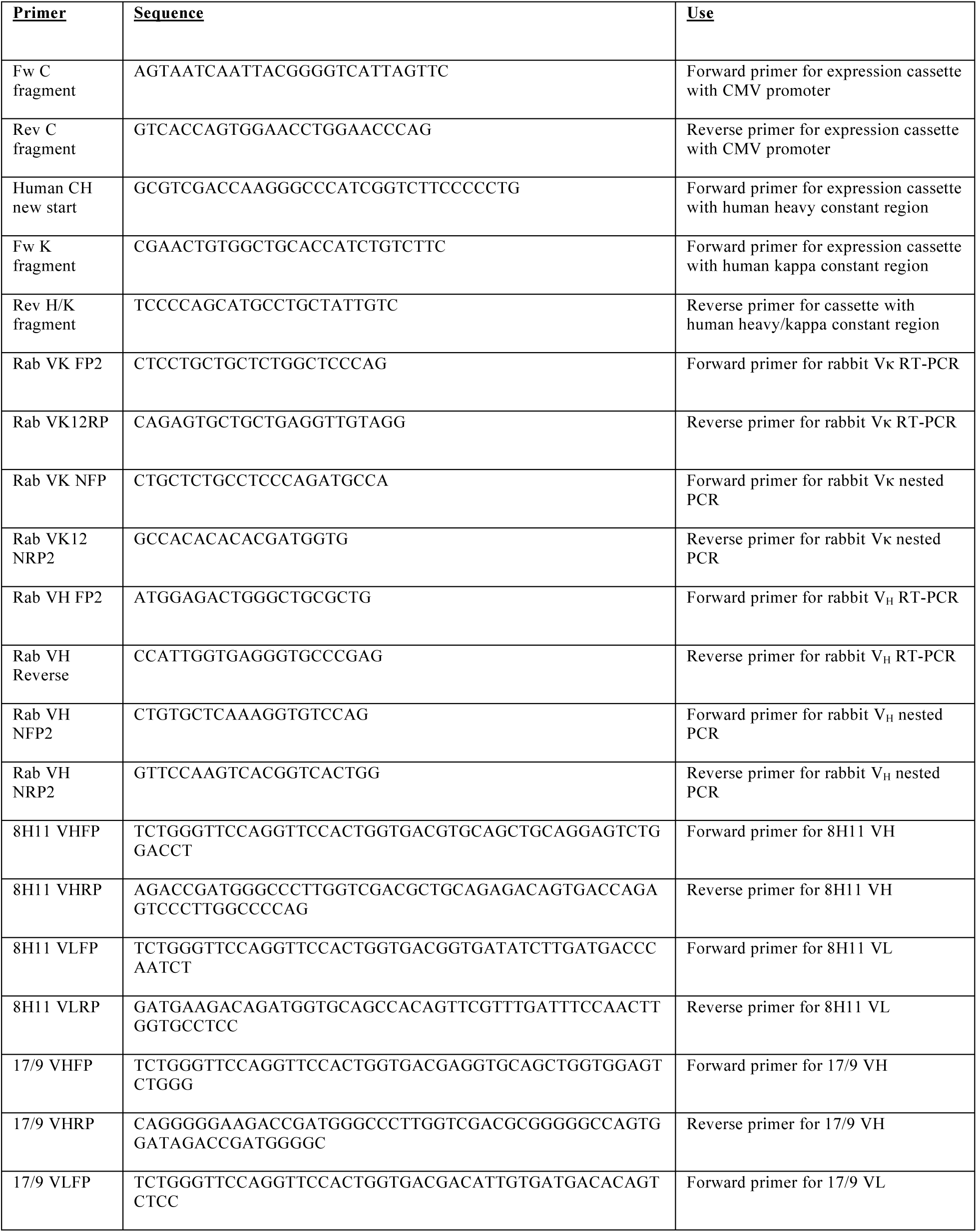
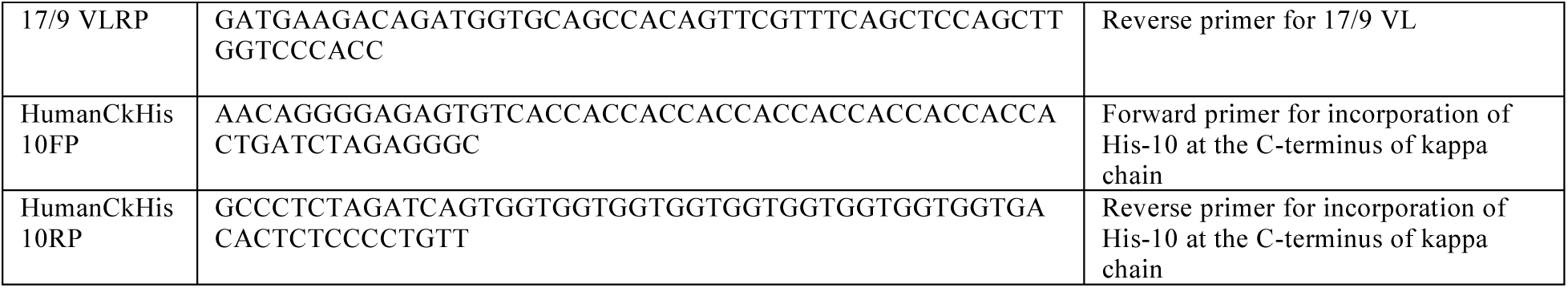
Primers used for rabbit single B cell cloning and antibody expression construct generation.

### Expression and purification of human chimeric antibodies

293T cells were co-transfected with plasmids encoding cognate heavy and light chains using the Xtreme Gene 9 DNA transfection reagent (Roche, Basel, Switzerland), then grown in Dulbecco’s modified Eagle medium supplemented with 10% fetal bovine serum (FBS, (Thermo Fisher Scientific, Waltham, MA, USA). Culture supernatants were collected 48 hours later for antibody screening, analysis, or purification. Ten-ml culture supernatants were concentrated to 1 ml on an Amicon ultrafiltration column (MilliporeSigma, Burlington, MA, USA; 50 kDa molecular weight cut off). Concentrated culture supernatants containing non-His-tagged mAbs were diluted 3-fold with Buffer A (0.02 M sodium dihydrogen phosphate, 0.15 M sodium chloride, pH 8.0). Protein A resin suspension (0.2 ml) (MilliporeSigma, Burlington, MA, USA) was equilibrated in Buffer A and incubated with the culture supernatant at 4°C for 1 hour with intermittent mixing. The resin was packed in a Poly-Prep chromatography column (Bio-Rad, Hercules, CA, USA) and washed with 20 volumes Buffer A. Antibody elution was performed in 0.1 M glycine-HCl buffer, pH 2.0, and neutralized by adding one-tenth volume 1M Tris-HCl, pH 9.0, to the eluted antibody fraction. Affinity chromatography was performed in a gravitational-flow mode with a ∼0.5 ml/min flow rate. The purified mAbs were dialyzed against phosphate-buffered saline (PBS, 0.1 M sodium phosphate, 0.15 M sodium chloride, pH 7.2) (Thermo Fisher Scientific, Waltham, MA, USA) containing 0.02% sodium azide in dialysis tubing with 6-8 kDa cutoff limit (Spectrum Chemical, New Brunswick, NJ, USA).

Concentrated culture supernatants containing His-tagged mAbs were diluted 3-fold with 50 mM Tris-HCl, pH 8, containing 0.1 M sodium chloride (HS-TBS; High-strength tris buffered saline) to which imidazole was added to a 5-mM final concentration (HS-TBS/5mM imidazole). Ni-NTA resin (0.3 ml) (Qiagen, Hilden, Germany) was pre-equilibrated with HS-TBS/5 mM imidazole and mixed with the diluted culture supernatant. The resin was incubated for 30 minutes at 4°C with intermittent mixing. The resin was then packed in a Poly-Prep chromatography column (Bio-Rad, Hercules, CA, USA) and washed with 30 column volumes of HS-TBS/5mM imidazole, followed by washing with 10 column volumes each of HS-TBS containing 10 mM and 20 mM imidazole. Stepwise elution of His-tagged mAbs was performed with two column volumes each of HS-TBS containing 50-, 100-, 200- and 300- mM imidazole. Chromatography was performed in a gravitational-flow mode with a ∼0.5 ml/min flow rate. Fractions were analyzed for the presence of mAb by 12% SDS-PAGE with silver staining. Fractions containing purified mAbs were pooled and dialyzed against PBS, pH 7.2, containing 0.02% sodium azide.

### Immunoblotting

Ten µl concentrated culture supernatant containing expressed human chimeric antibodies or Ni-NTA column flow-through was fractionated on 12% SDS-PAGE and electroblotted onto a polyvinylidene difluoride membrane (MilliporeSigma, Burlington, MA, USA). After overnight blocking at 4°C in 5% w/v non-fat dried milk in 1X TBS-T buffer (20 mM Tris pH 7.6, 160 mM sodium chloride, 0.05 % Tween 20), the blots were probed for the presence of heavy and light chain bands using either horseradish peroxidase (HRP)-conjugated polyclonal goat anti-human IgG Fc antibody or polyclonal goat anti-human kappa chain antibody (Thermo Fisher Scientific, Waltham, MA, USA). Antibody-coupled HRP activity was detected with Supersignal West Dura Extended Duration Substrate (Thermo Fisher Scientific, Waltham, MA, USA). Densitometric analysis of immunoblots was done using ImageJ software.

### Immunization of a New Zealand White (NZW) rabbit

A single, female NZW rabbit (Charles River, Saint-Constant, QC, Canada) was immunized intravenously with 0.2 ml PBS, pH 7.3 (without Calcium and Magnesium; 1X Gibco PBS), containing 5 x 10^11^ pfu (equivalent to 25 µg) CsCl-equilibrium-gradient-purified filamentous fd-phage (kindly provided by George P. Smith, Univ of Missouri-Columbia, USA) in the absence of adjuvant. Anti-phage antibody titer was measured bi-weekly, and a booster dose was performed at the end of week 7, once the primary antibody titer waned after reaching maximal value. The second immunization was administered both subcutaneously and intraperitoneally with 50 µg fd phage in 0.2 ml 1X Gibco PBS per site. The spleen was harvested 7 days following the booster injections and processed for B-cell sorting. All immunizations were carried out by staff in Simon Fraser University’s Animal Care Facility following the protocols reviewed and approved by the University’s Animal Care Committee (UACC).

### Preparing single-cell suspensions of rabbit splenocytes

The rabbit’s spleen was placed in a sterile tissue culture dish with pre-warmed 1X Gibco PBS, and after being dissected into small pieces, the tissue was dissociated by passage through a sterile, 20-gauge syringe. The tissue suspension was then passed through 40 µm cell strainer (VWR international, Randor, PA, USA) and the cell suspension was collected. Cells were harvested by centrifugation at 720g for 5 minutes. Red blood cells were lysed by gently resuspending the cell pellet in 9 ml 0.17 M ammonium chloride per gm cell pellet followed by incubation at room temperature for 5 minutes. Splenocytes were harvested and washed with 1X Gibco PBS and finally with pre-warmed RPMI 1640 media containing 10 % FBS. The cells were again washed with 1X Gibco PBS and counted in a hemocytometer upon removal of adherent cells. The cell density was adjusted to 5 x 10^6^ cells/ml; 5 ml cell suspension was layered over 5 ml Lympholyte^®^-Rabbit (Cedarlane Laboratories Inc., Burlington, Canada) and centrifuged at 1500g for 30 minutes at room temperature. The mononuclear-cell layer was aspirated, washed in RPMI 1640 media and counted, then pelleted and resuspended at a concentration of 6.5 million cells per ml in freezing media (90% FBS; 10% DMSO). Aliquots were frozen at −80°C overnight and later transferred to the vapor phase of a liquid nitrogen freezer.

### Preparation of antigen-specific reagents for labeling cells and for SPR analysis

The N-terminal 17 residues of the major coat protein, pVIII, of the filamentous bacteriophage, fd, comprises the majority of the surface of the virion, along with 4 minor coat proteins. A biotinylated peptide comprising this sequence (AEGDDPAKAAFDSLQAA-Orn (Biotin)-GC; Bio-pVIII peptide; GL Biochem Ltd., Shanghai, China) was complexed to APC-Cy7-labeled streptavidin (BD Biosciences, Franklin Lakes, NJ, USA) for antigen-specific labeling of immune, splenic B cells. Briefly, 4 µg (∼66 pmol) of APC-Cy7-streptavidin was incubated with a 50-fold molar excess of Bio-pVIII peptide in the presence of 1 mg/ml biotin-free bovine serum albumin (BSA) in 1X Gibco PBS in a total volume of 100 μl at 4°C overnight. Similarly, biotin-blocked PerCP-labeled streptavidin (BD Biosciences, Franklin Lakes, NJ, USA) was prepared by incubation with a 50-fold molar excess of biotin (MilliporeSigma, Burlington, MA, USA) under identical conditions. In addition, biotin-blocked, unlabeled streptavidin was prepared by incubating 190 µg streptavidin (Roche, Basel, Switzerland) in the presence of a 100-fold molar excess of biotin in 1X Gibco PBS in a final volume of 250 μl. Similar reactions were also set up for producing complexes of Bio-pVIII peptide and streptavidin in presence of 20-fold molar excess of biotin to ensure that no more than one Bio-pVIII peptide was bound per streptavidin tetramer, so as to act as a monovalent analyte in SPR analysis. Unbound Bio- pVIII peptide and biotin were removed by washing reaction mixes repeatedly on a 30 kDa cut-off ultrafiltration columns (MilliporeSigma, Burlington, MA, USA) at 4°C using a total of 2.5 ml 1X Gibco PBS. Concentrations of the protein complexes were quantitated by measuring absorbance at λ_280_.

### Sorting antigen-specific rabbit B cells

Frozen rabbit splenocytes were thawed rapidly at 37°C and were washed twice in 10 ml prewarmed RPMI 1640 media containing 10% FBS. The cells were then washed once in chilled staining buffer (1X Gibco PBS containing 10% FBS and 0.1% sodium azide). The cells were incubated with Fc blocking reagent (Innovex Biosciences, Richmond, Canada) following manufacturer’s recommendations. After washing, 20 µg biotin-blocked, unlabeled streptavidin were added to 10^6^ cells and incubated on ice for 15 minutes to block non-specific binding sites for streptavidin. Cells were then incubated with 0.3 µg of biotin-blocked PerCP-labeled streptavidin and 0.3 µg of APC-Cy7-labeled streptavidin complexed with Bio-pVIII peptide along with a 1:25-fold diluted FITC-labeled goat anti-rabbit IgG-Fc polyclonal antibody (Jackson ImmunoResearch, West Grove, PA, USA) in a final volume of 100 µl for 15 minutes at 4°C. Cells were then washed and resuspended in staining buffer for sorting.

The fluorophores used in sorting were spectrally well separated and there was no spectral overlap among the different detector channels used. Fluorescence-activated cell sorting (FACS) was performed on a BD FACS Jazz (BD Biosciences, Franklin Lakes, NJ, USA) using a 100 μm sort nozzle. Sorting was controlled by using BD FACS Sortware software (BD Biosciences, Franklin Lakes, NJ, USA). Rabbit mononuclear cells were identified by size and granularity using forward scatter area (FSC-A) *versus* side scatter area (SSC-A) plot. The antigen-specific cells were gated based on APC Cy7^+^ Bio-pVIII signal. FITC-IgG^+^ and biotin-blocked Per-CP streptavidin negative cells from this gate was sorted into wells containing 3 µl chilled lysis buffer (0.5X Gibco PBS, 3 U/µl RNaseOUT and 10 mM DTT) for a subsequent reverse-transcriptase reaction in a 96 well PCR plate. Post-sort analysis was performed using Flowing software (Turku Bioscience, Finland).

### Amplification of V_H_ and V_L_ coding sequences from single B cells

The 96-well PCR plate containing single antigen-specific B cells in 3 µl lysis buffer per well was briefly centrifuged after sorting, then mixed with 2 µl master mix (2.5% IGEPAL, 75 ng/µl random hexamers and 2.5 mM dNTPs). The resulting 5 µl mix was incubated at 65°C for 5 minutes and the reaction mix was chilled on ice where 5 µl reverse transcriptase mix (containing 2X RT buffer, 10 mM MgCl_2_, 14 mM DTT and 2 U/µl Superscript III RT enzyme) was added. The 10-µl reverse transcriptase reaction contained a final concentration of 1 U/µl RNaseOUT, 0.5% IGEPAL, 15 ng/µl random hexamer, 0.5 mM dNTPs and 1 U/µl Superscript III enzyme, and was performed at 42°C for 10 minutes, 25°C for 10 minutes, 50°C for 60 minutes, 85°C for 5 minutes and hold at 4°C. Three µl of reverse transcriptase reaction was used separately for light and heavy chain PCR amplification in a 25 µl reaction using Platinum Taq DNA polymerase. The PCR product (3 µl) from the first round of PCR was used as template for nested PCR reaction. The set of primers used in RT-PCR and nested PCR are listed in table 1. The PCR cycling conditions were: 40 cycles at 94°C for 2 minutes (initial denaturation), 94°C for 15 seconds (denaturation), 52°C or 50°C (annealing for RT-PCR or for nested PCR, respectively) for 30 seconds, 68°C for 1 minute (extension). All the reaction components were from Thermo Fisher Scientific except IGEPAL (MilliporeSigma, Burlington, MA, USA).

### Enzyme-linked immunosorbent assays (ELISAs)

The hemagglutinin (HA) peptide (YPYDVPDYAGAGC) or N-terminally biotinylated HA peptide (Bio-HA) (GL Biochem Ltd., Shanghai, China) or HER2 extracellular domain (ECD) (Acrobiosystems, Newark, DE, USA) was used as antigen in ELISAs. Wells of high binding-capacity microtiter plates (Corning Inc., Corning, NY, USA) were coated overnight at 4°C with 200 ng peptide or 1 µg protein (e.g., BSA, HER2 ECD or streptavidin) in 35 µl Tris buffered saline (TBS) (20 mM Tris-HCL, 150 mM sodium chloride, pH 7.5). Wells were blocked for 1 hour at 37°C in TBS containing 5% non-fat dried milk and 0.1% Tween-20. Streptavidin-coated wells were blocked with TBS containing 2% BSA, washed and incubated with 200 ng appropriate biotinylated peptide for 30 minutes at room temperature. After washing, wells were incubated for 2 hours at room temperature with mAbs as indicated. Bound mAb was detected using goat anti-human (H+L)-HRP (1:1000 dilution) (Thermo Fisher Scientific, Waltham, MA, USA). In the case of His-tag detection, a rabbit anti His-tag antibody (Proteintech, Rosemont, IL, USA) was used as primary antibody for detection of His-tagged proteins, whereas HRP-conjugated anti-rabbit IgG (Proteintech, Rosemont, IL, USA) was used to detect the rabbit primary antibody. Bound HRP was detected using 400 µg/ml ABTS (MilliporeSigma, Burlington, MA, USA) in citrate-phosphate buffer containing 0.03% v/v hydrogen peroxide. Each assay was performed in triplicate for statistical analysis. OD_405-490_ was measured using a Tecan M200 Pro microplate reader and data were plotted after subtraction of background signals from BSA-coated wells.

For competitive ELISAs, 1 nM (for chimeric 17/9 mAb) or 2 nM (for chimeric 8H11 mAb) non-His-tagged or His-10 human chimeric mAb was mixed with different concentrations (from 0.05 to 500 nM) of either HA peptide or HER2 ECD in PBS containing 1 mg/ml BSA and incubated at 4°C overnight to reach equilibrium. The mixture was then incubated with appropriate antigen immobilized in wells of high-capacity binding plates, and ELISAs were performed in triplicate as described. Data were fit to a 4-parameter logistic model using Microsoft Excel to calculate IC_50_ values [55].

### Determination of the molecular weights of native and recombinant mAbs using mass photometry

Mass photometry measurements were performed on a Two MP mass photometer (Refeyn Ltd.). Borosilicate microscope coverslips were cleaned sequentially with water, isopropanol and water, followed by drying under a flow of clean nitrogen. Silicone gaskets were placed on the coverslips to create wells for sample loading. The microscope was focused on the coverslip surface with 18 µl 1X Gibco PBS buffer in the gasket well. Samples were diluted 10-fold in the gasket well with 1X Gibco PBS buffer such that the final concentration of each biomolecule was ∼2 nM. Sample binding to the coverslip surface was monitored for 60 seconds using the software Acquire MP (Refeyn Ltd., Version 2.3.0). A molecular-weight standard curve was prepared using a mixture of standard proteins ranging in size from 56-670 kDa. The mass error observed in the molecular-weight standard curve was less than 5%. Data analysis was performed using Discover MP (Refeyn Ltd., version 2.3.0).

### Binding affinity measurement using bio-layer interferometry (BLI) and surface plasmon resonance (SPR)

#### A. Using BLI to measure the affinity of mouse mAb 8H11 or human chimeric mAb 8H11 for HER2 ECD

Mouse mAb 8H11 was obtained from Cancer Therapeutics, Inc. (Los Angeles, CA, USA). The native or chimeric 8H11 mAbs were biotinylated in a 20-fold molar excess of EZ-link NHS-PEG4-Biotin (Thermo Fisher Scientific, Waltham, MA, USA) following the manufacturer’s recommendations. Binding experiments were performed in PBS, containing 0.1% BSA using Octet Red 96 (Sartorius, Goettingen, Germany). Biotinylated mAbs were immobilized on streptavidin-coated biosensors (Sartorius, Goettingen, Germany) for 60 seconds at 50 µg/ml concentration. After a baseline step, varying concentrations (0.5-50 nM) of HER2 ECD analyte were used to obtain the binding rate. Following association, PBS containing 0.1% BSA was passed over the antigen-antibody complexes and the rate of dissociation was measured. Five separate binding reactions were performed. Binding affinities were calculated from association and dissociation rates determined with ForteBio Data analysis software v11.1.

#### B. Using SPR to measure the affinity of human chimeric mAb 238A1 for biotinylated pVIII peptide

SPR experiments were performed using a BIAcore X100 (GE Healthcare, Marlborough, MA, USA). Purified His-10 238A1 mAb was immobilized on Flow Cell 2 of a NTA sensor chip while His-10 human chimeric 17/9 mAb was immobilized on Flow Cell 1 of the same chip to serve as control surface. HBS-P+ buffer (10 mM HEPES pH 7.4, 0.15 M NaCl, 0.05% v/v P20 surfactant, (GE Healthcare, Marlborough, MA, USA) was used as the running buffer. Single-cycle kinetic analysis was performed by passing a range of streptavidin Bio-pVIII peptide complex concentrations (12.5 to 500 nM). Biotin-blocked streptavidin (500 nM) was used as a negative control analyte. Affinity was estimated following 120 seconds association and 240 seconds dissociation phase after each injection with a flow rate of 20 µl/min. The sensor surface in Flow Cell 2 was regenerated by passing over it 350 mM Na_2_EDTA, pH 8, after each cycle. Binding experiments were performed in triplicate. Double-referenced, background-subtracted, binding curves were analyzed using BIAevaluation software (GE Healthcare, Marlborough, MA, USA) following standard procedures. Kinetic parameters and estimated affinity values were determined from curve fitting using a single-cycle kinetics 1:1 binding model.

### Curve fitting and statistical analysis

Data plotting, standard deviation calculations and curve fittings were performed using Microsoft Excel 2007.

## Results

### A genetic system for cloning human chimeric mAbs

We wanted to establish a genetic system to convert single B-cell-derived variable coding sequences into human chimeric antibodies so that common, anti-human secondary reagents could be used to perform downstream analysis. In addition, our interest was to develop an antibody purification or enrichment system that is simple, generic, specific, economical, and flexible for both small- and large-scale purification and can be used at high-throughput level.

To meet our objective, we used an antibody-gene assembly system adapted from Liao *et al*. [28]. This assembly system requires size-overlap extension PCR of three overlapping linear expression cassettes to generate the coding region of full-length chimeric mAbs. One of the expression cassettes (P fragment) has the P_CMV_ region and an immunoglobulin leader sequence [56]. Another common expression cassette (C fragment) used in this system encodes either the human kappa light chain constant region or the human gamma 1 heavy chain constant region (called C_κ_ or C_H_ fragment, respectively) (Fig 1A and C). However, the heavy chain C fragment cassette obtained from Liao *et al.* failed to express functional mAbs in our hands. Sequencing data revealed that the heavy chain C fragment from Liao *et al.* was missing 12 amino acids from the N-terminal region. We, therefore, engineered the heavy chain C-fragment cassette to include the missing amino acids (by using the corresponding sequence derived from plasmid 10E8 mAb heavy chain expression vector; see *Material and methods*) which ultimately resulted in generation of a functional expression system that consistently produced chimeric mAbs in mammalian cell culture. Furthermore, we constructed different sets of C fragments whose coding sequence for the human kappa or gamma-1 chain constant regions is joined to either a His-6 or His-10 tag at the C-terminal end, to explore the use of IMAC for antibody purification (Fig 1B and D). The coding sequence of each human constant region within a C fragment is followed by a stop codon and a poly A signal sequence. A third cassette (V fragment) comprises a user defined V_H_ or V_L_ sequence with overlapping leader and respective heavy/light chain constant domain flanking sequences. Size-overlap extension PCR is then used to assemble the three cassettes into a single human chimeric antibody heavy or light chain gene with or without a His-tag (Fig 1A-D).

### Cloning, expression, and purification of recombinant His-tagged human chimeric mAbs from mammalian cell culture

Different lengths and locations of polyhistidine tags were screened to optimize reliable and reproducible purification of recombinant chimeric mAbs from mammalian cell culture using IMAC. We used the strategy described above to generate a recombinant mouse-human chimeric form of a well-characterized mouse anti-HER2 8H11 mAb [57] containing a 6X or a 10X polyhistidine tag (His-6 or His-10, respectively) at the C-terminus of either the human kappa light or gamma-1 heavy chain. The incorporation of His-6 or His-10 tags at the C-terminus of the heavy chain is likely to impact effector functions of chimeric mAbs due to direct involvement of Fc regions in antibody effector functions [58, 59]. We, therefore, first attempted to purify 8H11 chimeric mAb derived from expression constructs bearing His-6 or His-10 fused to the C-terminus of the kappa light chain. To this end, we generated a blunt-end mouse-human chimeric 8H11 mAb heavy chain expressing gene and a kappa light chain gene terminated with a His-6- or His-10-encoding region. Each gene was ligated separately into the blunt end site of the pJET1.2/blunt cloning vector. The chimeric 8H11 heavy chain and one of the chimeric light chain clones (terminated with either His-6 or His-10) were used to co-transfect 293T cells for expression and secretion of chimeric recombinant 8H11 mAb into cell culture media. The corresponding non-His-tagged form of chimeric 8H11 light chain gene was also cloned and the 8H11 mAb devoid of any His-tag was expressed. The culture media, upon clarification, was used for purification of His-tagged chimeric 8H11 mAb using the Ni-NTA resin, whereas the non-His-tagged 8H11 mAb was purified using the Protein A resin. A densitometric analysis of immunoblots showing heavy and light chains of chimeric mAbs in the culture supernatant (Fig 4 A and B; *lanes 2-5*) suggests the expression levels of different chimeric mAbs were similar, with the difference in their expression being less than 1.5-fold.

Our attempts to IMAC-purify mAb 8H11 containing a His-6 at the C-terminus of the kappa light chain were unsuccessful, as an extremely poor yield was obtained (Fig 2A). Importantly, the incorporation of the more extended His-tag, His-10, at the C-terminus of the kappa light chain resulted in a large increase in the yield of the purified recombinant mAb (Fig 2B). The imidazole-eluted fractions from the column showed the presence of two major bands corresponding to the heavy and light antibody chains (Fig 2 B; *lanes 2-3)*. We also purified a non-His-tagged version of the chimeric 8H11 mAb to homogeneity using Protein A affinity chromatography (Fig 2C; *lane 1*). This non-His-tagged chimeric 8H11 light chain polypeptide clearly showed slightly increased mobility on SDS-PAGE gel as compared to its counterpart His-10 version (Fig 2C; *lanes 1 and 2*), whereas the mobility of the heavy chain bands in these chimeric 8H11 versions were identical. These antibodies were also analyzed under non-reducing conditions to confirm that the His-tagged light chain, observed under reducing conditions, was part of the full-length chimeric mAb (Fig 2D; *lanes 2 and 3*). These observations suggest that the presence of His-10 at the kappa chain C-terminus facilitates one-step enrichment of secreted recombinant chimeric 8H11 mAb directly from mammalian cell culture media. Typical mAb yields are about 2 μg/ml of culture supernatant for non-His- tagged recombinant antibodies and about 3 μg/ml of culture supernatant for His-10 recombinant antibodies. In contrast, incorporation of His-6 or His-10 at the C-terminus of the heavy chain failed to facilitate purification of chimeric mAb (Supplementary Fig 1 and Fig 2E, respectively). The inefficiency of binding of mAbs with His-10 heavy chain or His-6 kappa chain to the Ni-NTA resin accounts for some of these observations (Fig 4A and B; *lanes 7-8*).

**Figure 2.**
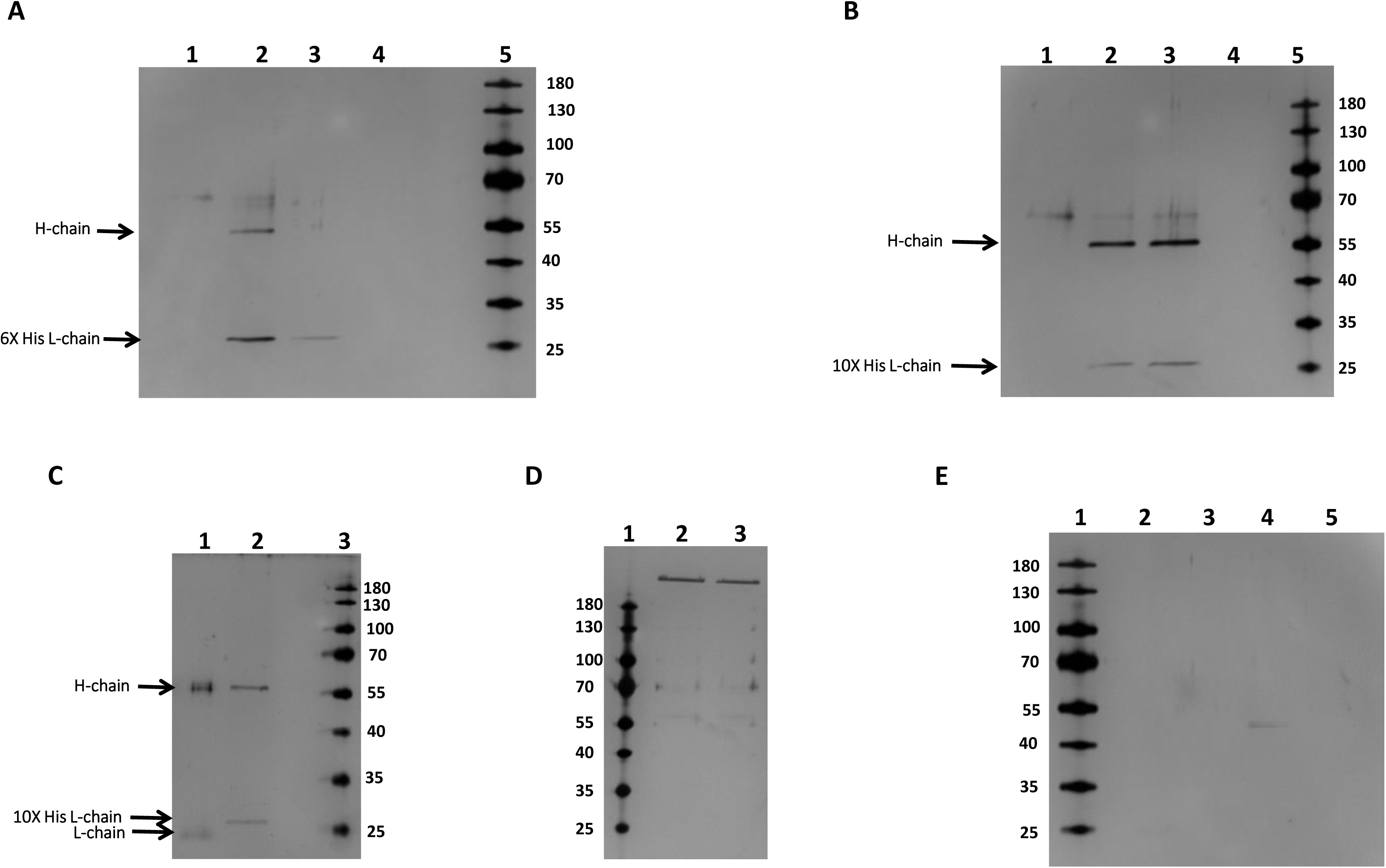
SDS-PAGE showing the purified His-tagged and non-His-tagged forms of mouse-human chimeric 8H11 mAb: Specific heavy chain (referred as H-chain) and light chain (referred as L-chain) polypeptides were observed to migrate with an apparent molecular weight of ∼ 55 kDa and ∼27 kDa in the SDS-PAGE gels. The numbers indicated adjacent to each gel represent the molecular weight of polypeptides (in kDa) in the protein ladder. (A) Polyacrylamide gel showing the different eluent fractions (50-, 100-, 200- and 300-mM imidazole) from Ni-NTA column derived from cell culture media expressing His-6 kappa chain containing human chimeric 8H11 mAb (*lanes 1-4*). (B) Different eluent fractions of His-10 kappa chain containing chimeric 8H11 mAb derived from the affinity column are shown (*lanes 1-4* represent 50-, 100-, 200-, and 300- mM imidazole fractions). (C) The gel represents pure fractions of human chimeric 8H11 proteins in the absence (*lane 1*) or presence (*lane 2*) of His-10, electrophoresed under reducing conditions. (D) The purified fractions of both non-His-tagged (*lane 2*) and His-10 forms (*lane 3*) of chimeric 8H11 mAbs are shown on a non-reducing SDS-PAGE gel. (E) Different imidazole fractions obtained from affinity chromatography (from 50- to 300- mM in *lanes 2-5*) of human chimeric 8H11 mAb with C-terminal His-10 IgG1 heavy chain are shown.

To ensure the generic nature of our purification scheme using His-10 at the C-terminus of chimeric mAb light chain, we cloned, expressed, and purified recombinant human chimeric version of another well-characterized mouse anti-hemagglutinin (anti-HA) 17/9 mAb, whose V_H_ and V_L_ coding sequences have been described [60, 61]. Once again, incorporation of His-10 at the C-terminal end of the light chain facilitated single-step purification of the chimeric mAb (Fig 3A, *lane 2*). To perform comparative activity analysis, we also purified a non-His-tagged version of the chimeric 17/9 mAb using Protein A affinity chromatography (Fig 3A; *lanes 3*). These mAbs were also purified as full-length chimeric mAbs as made evident by the absence of any free light chain from the non-reducing gel shown in Fig 3B.

**Figure 3.**
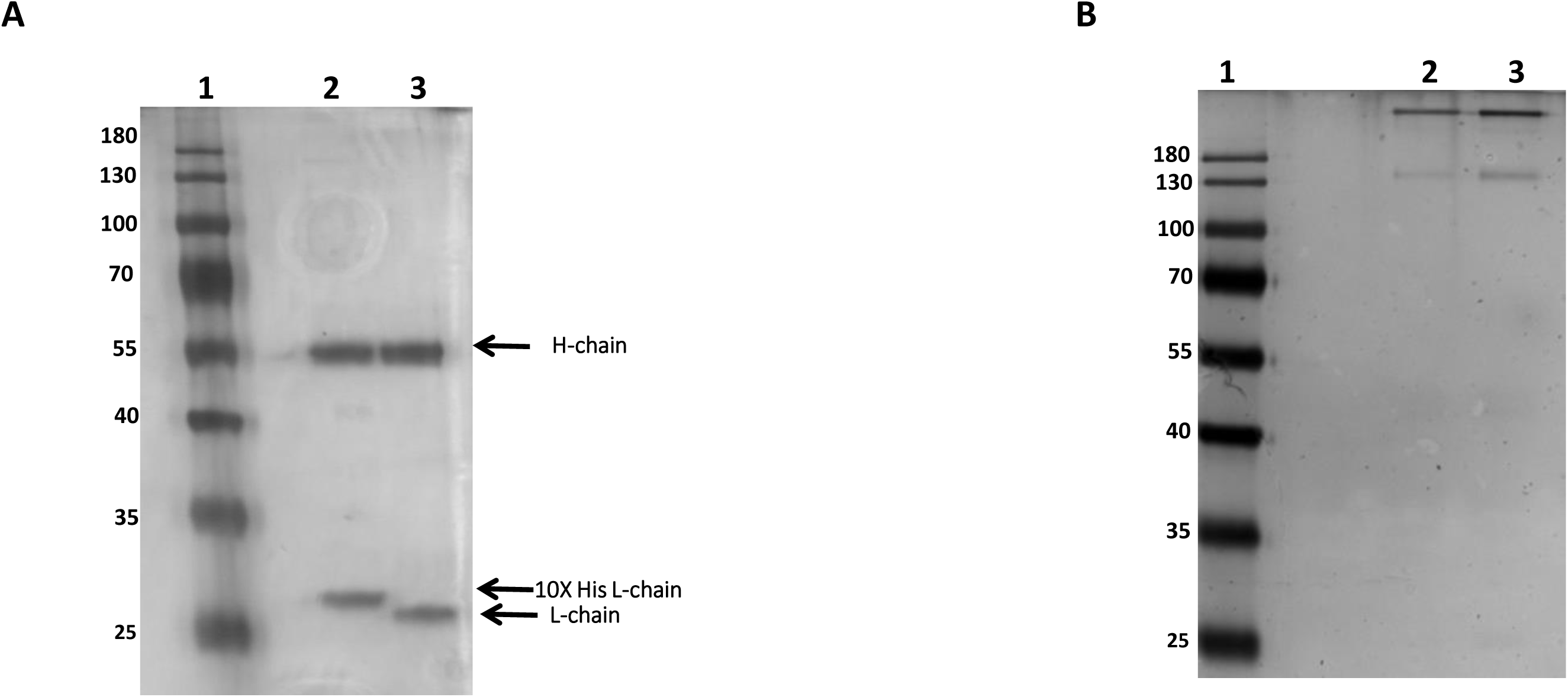
SDS-PAGE exhibiting the purified His-10 and non-His-tagged forms of mouse-human chimeric 17/9 mAb: (A) The purified fraction of His-10 kappa chain containing chimeric 17/9 mAb purified using Ni-NTA column (*lane 2*) and the purified form of non-His-tagged chimeric 17/9 mAb (*lane 3*) eluted from Protein A column are shown on a reducing SDS-PAGE gel. Specific light chain (referred to as L-chain) and IgG1 heavy chain (referred to as H-chain) polypeptides were observed as indicated by arrows. (B) The purified fractions of both His10 (*lane 2*) and non-His-tagged forms (*lane 3*) of chimeric 17/9 mAbs are shown on a non-reducing SDS-PAGE gel. The numbers to the left indicate the molecular weight of polypeptides (in kDa) in the protein ladder.

### Immunoblot analysis for estimating the expression levels of various recombinant chimeric mAbs

Our inability to purify recombinant chimeric mAbs upon incorporation of His-6 or His-10 at the C-terminus of the chimeric heavy chain or His-6 at the C-terminus of the chimeric light chain raised an important question as to whether these observations are largely due to poor expression of these chimeric constructs, or to the inability of expressed recombinant mAbs to bind to IMAC resin. We performed immunoblot analysis on supernatants from cell cultures expressing recombinant chimeric 8H11 mAb containing different lengths and locations of His-tag. These immunoblots were developed using either anti-human IgG-Fc or anti-human kappa chain secondary antibodies. Our data suggest that incorporation of His-tags at the C-terminus of either human kappa light chain or human heavy chain did not significantly inhibit the expression of chimeric 8H11 light chain or heavy chain; the difference between their expression levels was not more than 1.5-fold as measured by densitometric analysis (Fig 4 A and B; *lanes 2-5*). However, immunoblot analysis of the column flow-through, obtained after the chimeric mAbs were allowed to bind to Ni-NTA resin, showed that only the recombinant mAb bearing His-10 fused to the kappa chain bound efficiently to the resin, as no unbound light or heavy chain was observed in the flow-through (Fig 4 A and B; *lane 9*). In contrast, the recombinant 8H11 mAb bearing either His-6 at the C-terminus of the kappa light chain or His-10 at the C-terminus of the heavy chain failed to bind strongly to the resin, as was evident from the appearance of the corresponding light and heavy chains in the column flow-through (Fig 4 A and B; *lanes 7 and 8*). Taken together, these observations show that the presence of His-10 at the C-terminus of the kappa chain allowed the chimeric mAb to bind to the Ni-NTA resin, resulting in significantly higher yields upon IMAC purification.

**Figure 4.**
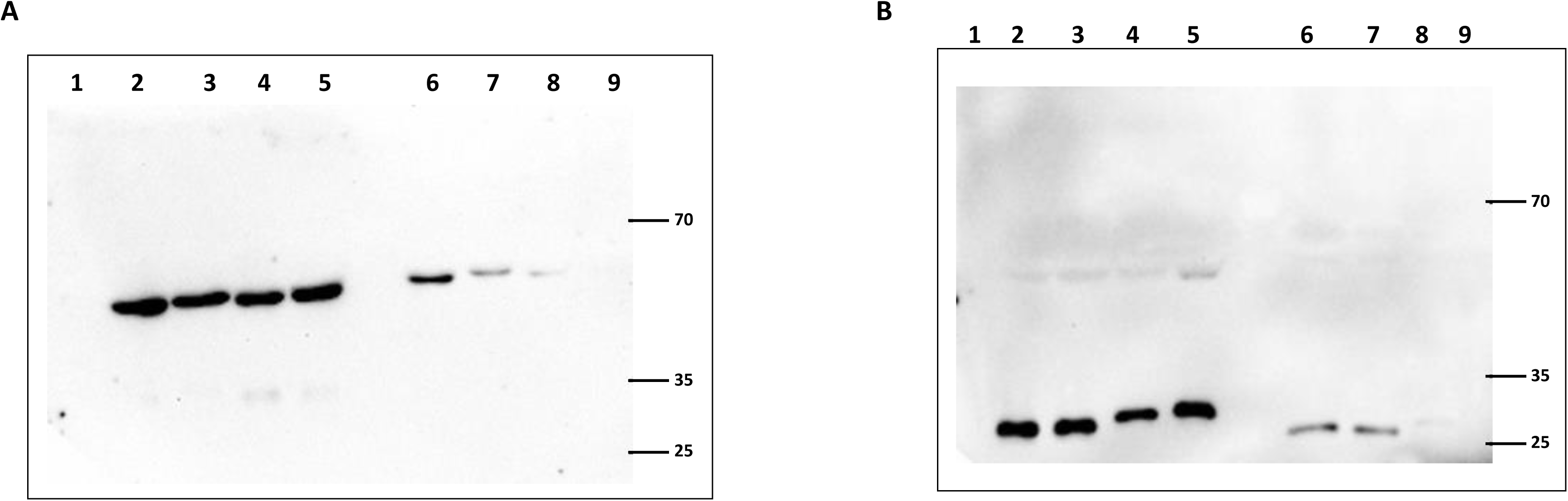
Immunoblots for analyzing the expression level and affinity resin binding capacity of different chimeric 8H11 antibodies with or without His-tag: (A) *Lanes 1-5* represent immunoblot analysis of human chimeric heavy chain in cell culture supernatant derived from 293T cells expressing different 8H11 heavy and light chain pairs. *Lane 1*: Negative control supernatant; *Lane 2*: Chimeric 8H11 without His-tag; *Lane 3*: Chimeric 8H11 with His-10 heavy chain; *Lane 4*: Chimeric 8H11 with His-6 light chain; *Lane 5*: Chimeric 8H11 with His-10 light chain. *Lanes 6-9* represent immunoblot analysis of human chimeric 8H11 heavy chain in Ni-NTA column flow-through after incubation of 3-fold dilute cell culture supernatant with Ni-NTA resin. *Lane 6*: Chimeric 8H11 without His-tag; *Lane 7*: Chimeric 8H11 with His-10 heavy chain; *Lane 8*: Chimeric 8H11 with His-6 light chain; *Lane 9*: Chimeric 8H11 with His-10 light chain. (B) The immunoblot analysis shows human chimeric 8H11 light chain in different culture supernatant and column flow-through in the same lane order as described for figure 4A. The numbers to the right indicate the molecular weight of polypeptides (in kDa) in the protein ladder.

### Effect of chimerization and incorporation of His-10 on the binding affinity of chimeric mAbs

Following purification of recombinant non-His-tagged and His-10 chimeric mAbs, we performed a preliminary analysis to determine whether these chimeric forms show antigen binding capacity. To this end, both the recombinant chimeric forms of 8H11 mAb were analyzed in an indirect ELISA to confirm binding to immobilized human HER2 ECD. Detection of binding signal by anti-His antibody was dependent on the presence of His-10 in the light chain of chimeric 8H11 mAb. This confirmed that the His-10 chimeric 8H11 mAb retains the capacity to bind to its specific antigen, and that the His-10 tag in the kappa light chain is accessible to the anti-His antibody (Fig 5A). As expected, non-His-tagged chimeric 8H11 mAb failed to show the antigen binding signal when probed with the anti-His antibody, whereas the same chimeric antibody showed HER2 ECD binding activity when probed with an HRP-conjugated anti-human secondary antibody (Fig 5A; *Gray bar*). These observations suggest that chimerization and the addition of His-10 to the C-terminus of the kappa light chain does not affect the antigen-binding activity of the chimeric 8H11 mAb.

**Figure 5.**
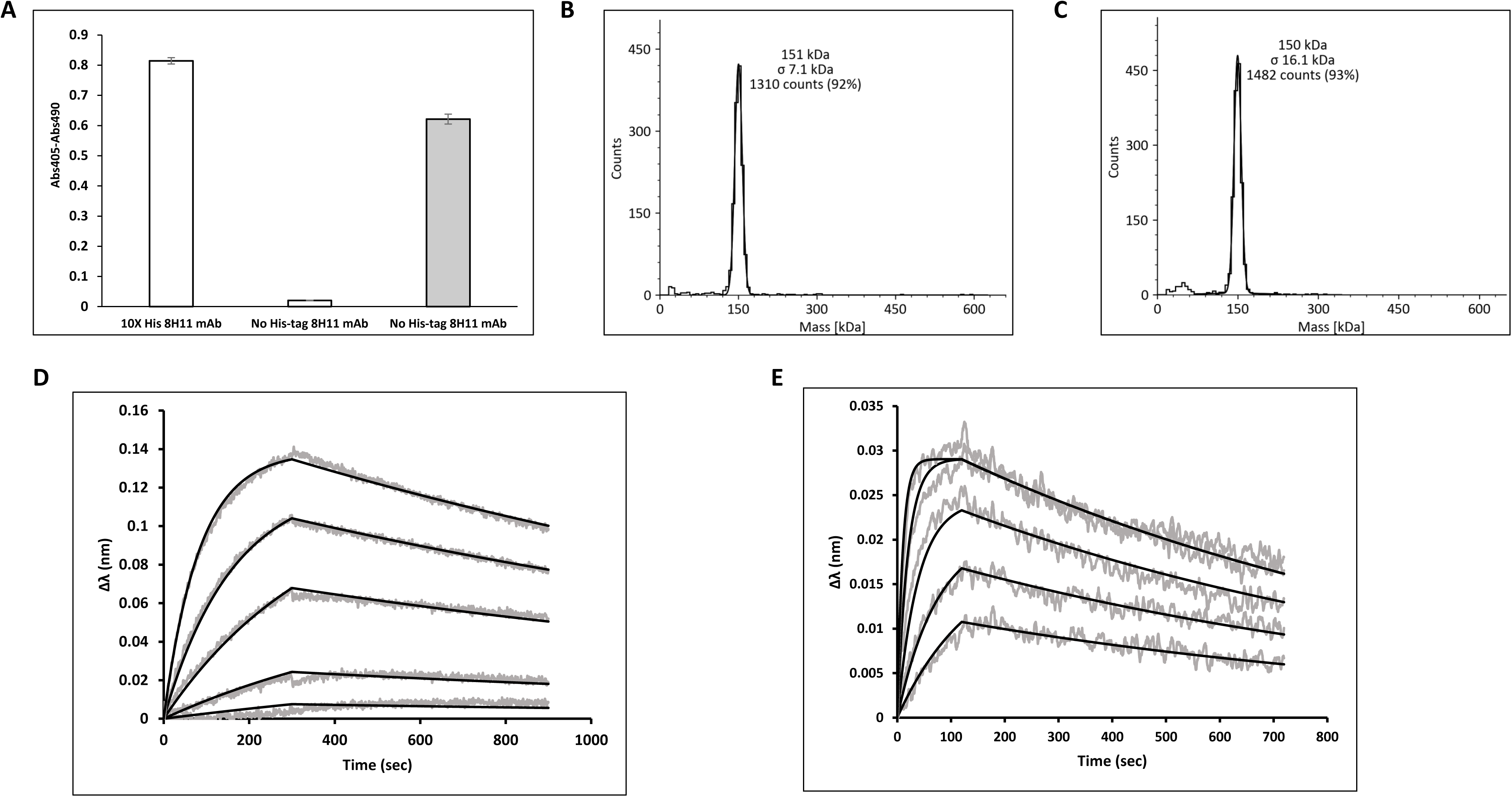
Analysis of binding of native mouse 8H11 and recombinant human chimeric 8H11 mAb to their antigen HER2 ECD: (A) The bar diagram represents the binding of purified His-10 or non-His-tagged 8H11 chimeric mAbs in ELISA. ELISA wells coated with HER2 ECD were incubated with purified His-10 or non-His-tagged chimeric 8H11 mAb as indicated. The presence of His-10 in the bound mAb was identified using a rabbit anti-His antibody. HRP conjugated anti-rabbit IgG was used to indirectly detect the presence of His-10 in the primary antibody (*White bars*). The non-His-tagged human chimeric mAb binding to the antigen was also detected using HRP-conjugated anti-human IgG (H+L) in parallel binding reactions. (*Gray bar*). The values represent the mean optical density obtained from triplicate experiments and error bars represent the standard deviations. (B and C) The histogram represents the observed molecular mass distribution of mouse 8H11 (B) and chimeric non-His-tagged form of 8H11 mAb (C) in solution as measured by mass photometer. (D and E) Binding of HER2 ECD to biotinylated mouse mAb 8H11 (D) or to biotinylated recombinant human chimeric non-His-tagged 8H11 (E) immobilized on streptavidin biosensor. Affinity measurements were conducted using BLI in 5 separate binding reactions.

Next, we wanted to analyze the effect of chimerization on the binding affinity of 8H11 mAb. We performed an initial analysis to determine the molecular weight of the native and chimeric 8H11 mAbs using mass photometer analysis. This technique allows the estimation of molecular weight of biomolecules in solution by analyzing the backscattered light in a microscope. Both native and chimeric forms of 8H11 exhibited a single dominant peak at 151 ± 7.1 kDa and 150 ± 16.1 kDa, respectively (Fig 5B and C). These observations suggest that purified forms of both native and chimeric 8H11 exist as monomeric species in solution and that chimerization did not affect the monomeric status of the mAb.

To evaluate the effect of chimerization on binding affinities, we performed comparative binding analysis of native mouse 8H11 mAb and non-His-tagged human chimeric 8H11 recombinant mAb. The 8H11 mAb was chosen for this analysis due to the availability of both fully murine and mouse-human chimeric forms of this mAb. Both mAbs were biotinylated and immobilized on streptavidin biosensors on an octet BLI system. Binding affinities were then measured in different concentrations of human HER2 ECD as analyte. The association and dissociation curves were fitted to a 1:1 binding model, which yielded dissociation constants (K_d_s) of 2.14 ± 0.02 nM for the interaction between mouse 8H11 and HER2 ECD and 3.57 ± 0.08 nM for non-His-tagged human chimeric 8H11 and HER2 ECD (Fig 5D and E). Thus, mouse-human IgG1 chimerization resulted in a ∼40% reduction in the binding affinity of mouse 8H11 mAb.

It is also important to analyze the effect of addition of His-10 on binding affinity of the chimeric antibody. To this end, we compared the relative affinity of both non-His-tagged and His-10 forms of the purified chimeric 8H11 mAb in a competitive ELISA. Both forms of chimeric 8H11 mAbs were separately allowed to bind to gradually increasing concentrations of human HER2 ECD antigen in solution and the equilibrium mixture was added to the surface immobilized antigen in an ELISA plate. The resulting binding signal was obtained, and 4 parameter logistic model was used to fit the curve as shown in Figure 6A. Both forms of the chimeric 8H11 antibody exhibited overlapping binding curves with IC50 values of 1.87 nM and 1.49 nM for His-10 and non-His-tagged forms, respectively (Fig 6A). These data, therefore, indicate that incorporation of His-10 at C-terminus of the kappa light chain did not strongly affect binding affinity of the chimeric antibody. To further validate these findings, we also performed a competitive ELISA with His-10 and non-His-tagged versions of human chimeric 17/9 mAb with HA peptide antigen. The corresponding IC50 values for His-10 and non-His-tagged forms of chimeric 17/9 mAb were 79.1 nM and 61.5 nM, respectively (Fig 6B). These data also indicate that adding His-10 tag using our expression cassette did not strongly affect binding affinity of chimeric mAbs.

**Figure 6.**
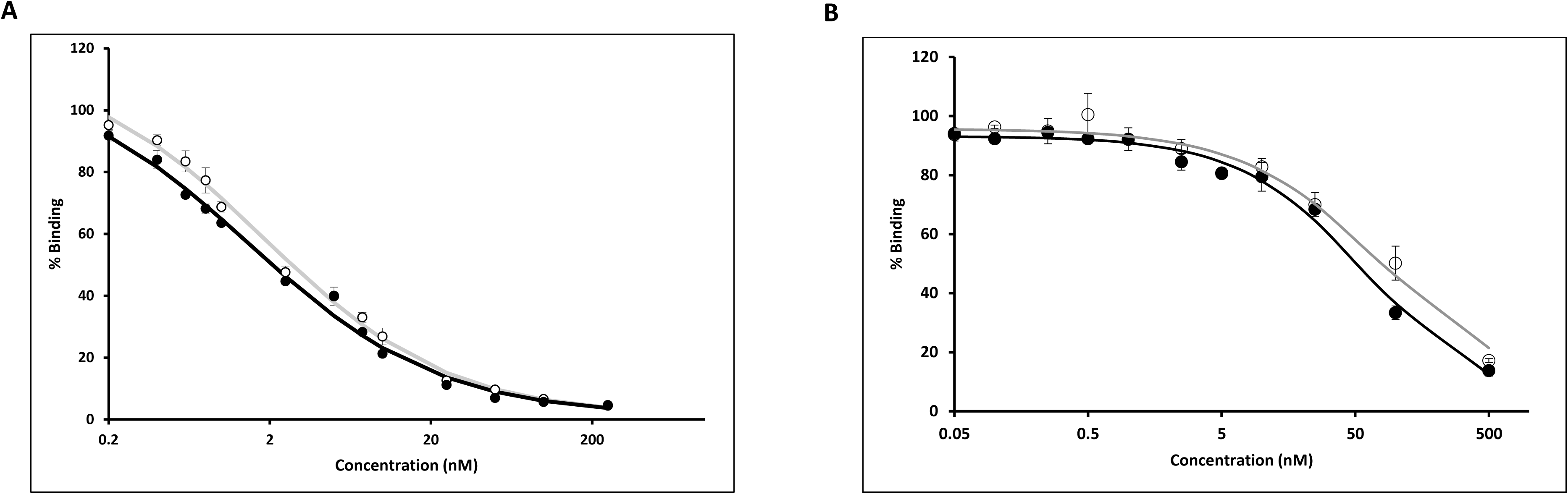
Competitive ELISA of His-10 and non-His-tagged forms of (A) chimeric 8H11 mAb or (B) chimeric 17/9 mAb: Purified His-10 (*open circles*) or non-His-tagged chimeric mAb (*filled circles*) was incubated with increasing concentrations of HER2 ECD or HA peptide, respectively, in solution and the equilibrium mix was added to wells coated with the corresponding antigen to measure the resulting free mAb. The % binding is indicated on the vertical axis considering the absorbance obtained in absence of antigen in equilibrium solution as 100%. The antigen concentration on the horizontal axis is indicated in logarithmic scale. The values represent the mean % binding obtained from triplicate experiments and error bars represent the standard deviations.

### Isolation of B cells specific for the major coat protein (pVIII) of fd bacteriophage

Following the establishment of a His-10 based purification system, we performed a proof-of-concept study to analyze the ease of integration of our cloning and purification system into a single B cell-based antibody discovery workflow. We chose to use pVIII, the most abundant coat protein of the filamentous bacteriophage, fd, as immunogen for our prototype study. To this end, we immunized a NZW rabbit with fd phage and analyzed serum antibody titers against the pVIII peptide before and after the booster dose to confirm a secondary immune response with higher magnitude. Splenic B lymphocytes were then isolated and labeled with complexes of APC Cy7-labeled streptavidin and Bio-pVIII peptide and with FITC-labeled anti-rabbit IgG. It is important to note that, in the absence of a general B-cell marker in rabbits, the antigen-specific B-cell labeling was restricted to anti-rabbit IgG (Fig 7). A small number of single, antigen-specific class-switched IgG^+^ B cells were sorted into wells of a 96-well plate containing lysis buffer.

**Figure 7.**
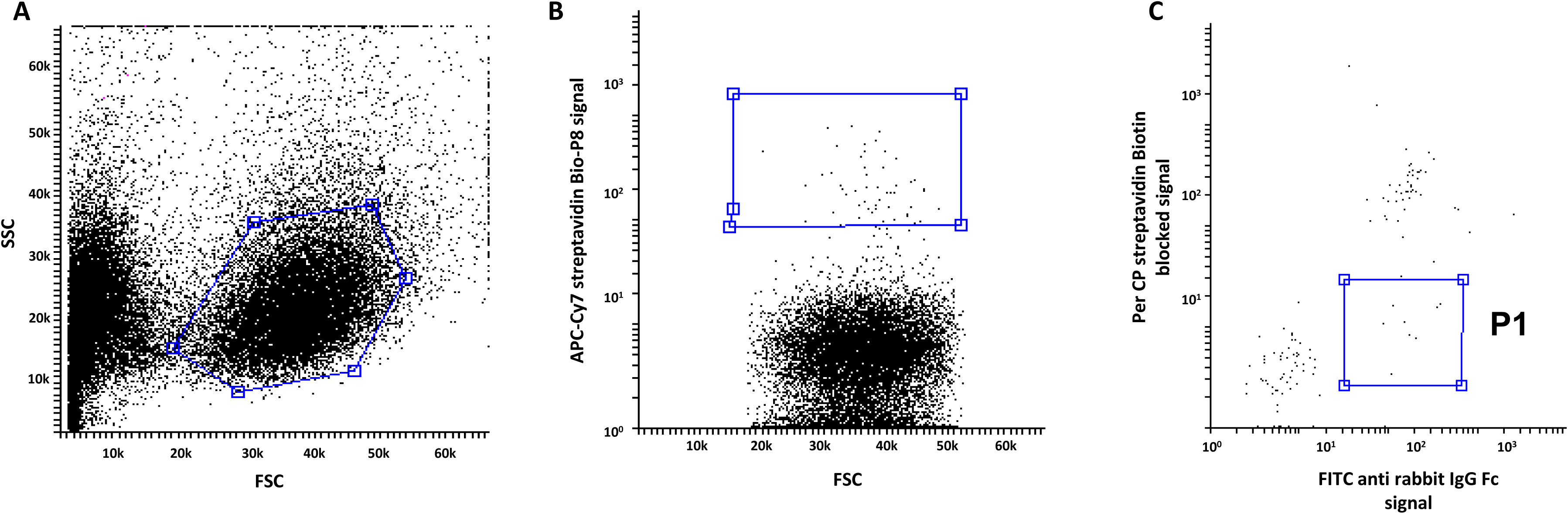
Hierarchical FACS gating strategy for isolation of antigen-specific B cells against the filamentous phage major coat protein, pVIII: Splenocytes were partially purified from the spleen of a rabbit after immunization with filamentous fd bacteriophage, and stained for FACS analysis and sorting of single, antigen-specific B cells. The hierarchical gates in dot plots (shown as boxes) were set as follows: (A) heterogeneous spleen mononuclear cells (B) APC Cy7^+^ labeled streptavidin-Bio-pVIII peptide complex binding cells (C) Biotin blocked Per CP-labeled streptavidin (decoy) negative and FITC IgG^+^ cells. Single cells from Gate P1 were sorted into a 96 well plate for single cell RT-PCR. The axis for forward and side scatter are in linear scale while those for the fluorescence signals are in logarithmic scale.

### Cloning, expression, and analysis of rabbit-human chimeric anti-pVIII mAbs

The naturally paired V_H_ and V_L_ gene segments from single antigen-specific B cells were amplified by RT-PCR followed by nested PCR reactions. This study used single forward and reverse primers for the PCR reactions to amplify rabbit V_H_ and V_L_ genes. The forward primer used in this study corresponds to conserved regions in the known rabbit antibody heavy or light chain leader gene sequences as derived from the IMGT IG germline gene database [62], whereas the reverse primer was derived from the constant region of either the IgG1 heavy or the kappa light chain coding region in rabbits (κ chains constitute the majority of light chains in the rabbit antibody repertoire). As an example, paired V_H_ and V_L_ gene segments were amplified from 2 of 5 single, antigen-specific IgG^+^ B cells that were sorted in a single experiment (Fig 8A and B). The amplified V_H_ and V_L_ gene segments derived from single B cells were converted into full-length rabbit-human chimeric antibody heavy chain genes and His-10 containing light chain genes using the tripartite linear expression cassette system depicted in Figure 1. The chimeric genes were separately cloned into the plasmid vector pJET 1.2.

**Figure 8.**
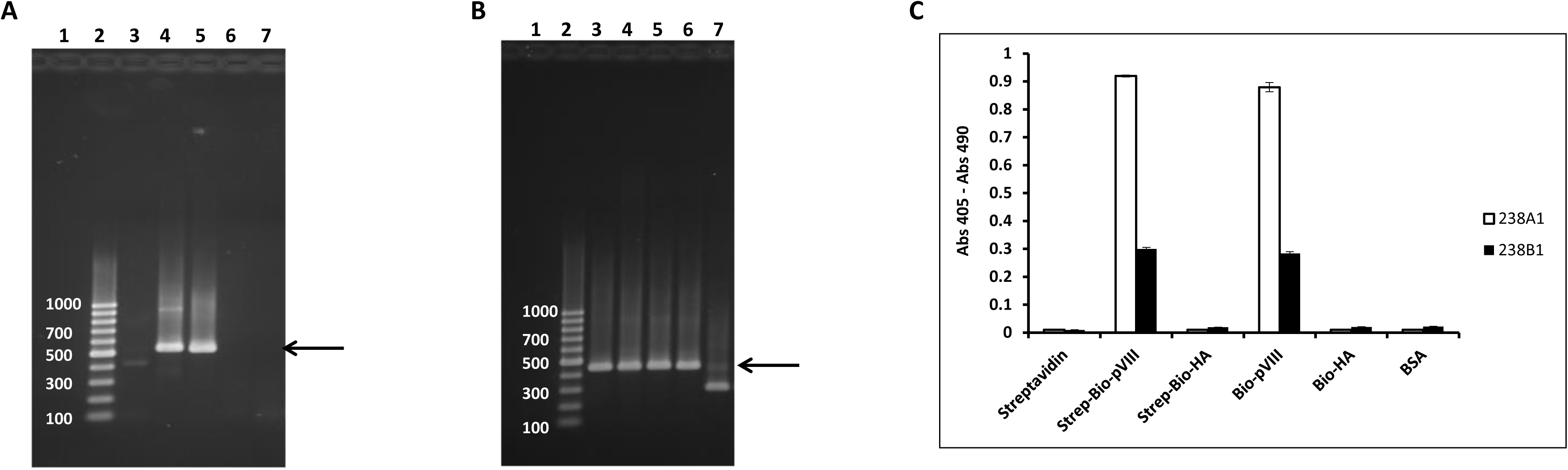
Isolation of antibody genes using antigen-specific B cells and analysis of binding of anti-pVIII mAbs: Representative agarose gels show RT-PCR products of V_H_ coding region (A) or V_L_ coding region (B) from 5 single, antigen-specific rabbit B cells. *Lane 1* in both the gels represents RT-PCR samples from “no cell” negative control. Desired product size is indicated by arrows alongside the 100-bp ladder. (C) Indirect ELISA was done to evaluate the specificity of binding of anti-pVIII antibodies (238A1 and 238B1). Wells of an ELISA plate was coated with peptides, proteins, or peptide-protein complexes, as indicated, and incubated with purified His-10 anti-pVIII rabbit-human chimeric mAbs. Bound mAb was identified using HRP-conjugated anti-human IgG antibody; optical densities (405 nm – 490 nm) are indicated on the vertical axis. The signal arising from 238A1 mAb binding is represented as *white bars* whereas the signal derived from 238B1 mAb binding is shown with *black bars*. The values represent the mean optical density obtained from triplicate experiments and error bars represent the standard deviations.

The adherent cell line, 293T, was transiently co-transfected with 2 plasmids expressing the V_H_ and V_L_ domains cloned from a single B cell. Culture media containing chimeric antibodies were harvested 60 h post-transfection and tested for antigen-specific binding in indirect ELISAs. Two single-cell derived, newly discovered antibodies (238A1 and 238B1) bound specifically to the pVIII peptide and did not bind to a non-specific peptide (HA peptide) or to unrelated proteins (streptavidin and BSA) (Fig 8C). The 238A1 mAb was chosen for further purification and characterization, as it produced a ∼3-fold stronger signal compared to mAb 238B1 (Fig 8C).

### SPR measurement of binding affinity of anti-pVIII chimeric mAb 238A1

The chimeric His-10 tagged 238A1 mAb was purified using IMAC (Fig 9A) and the purified fraction was used to estimate its molecular weight in solution using the mass photometer.

**Figure 9.**
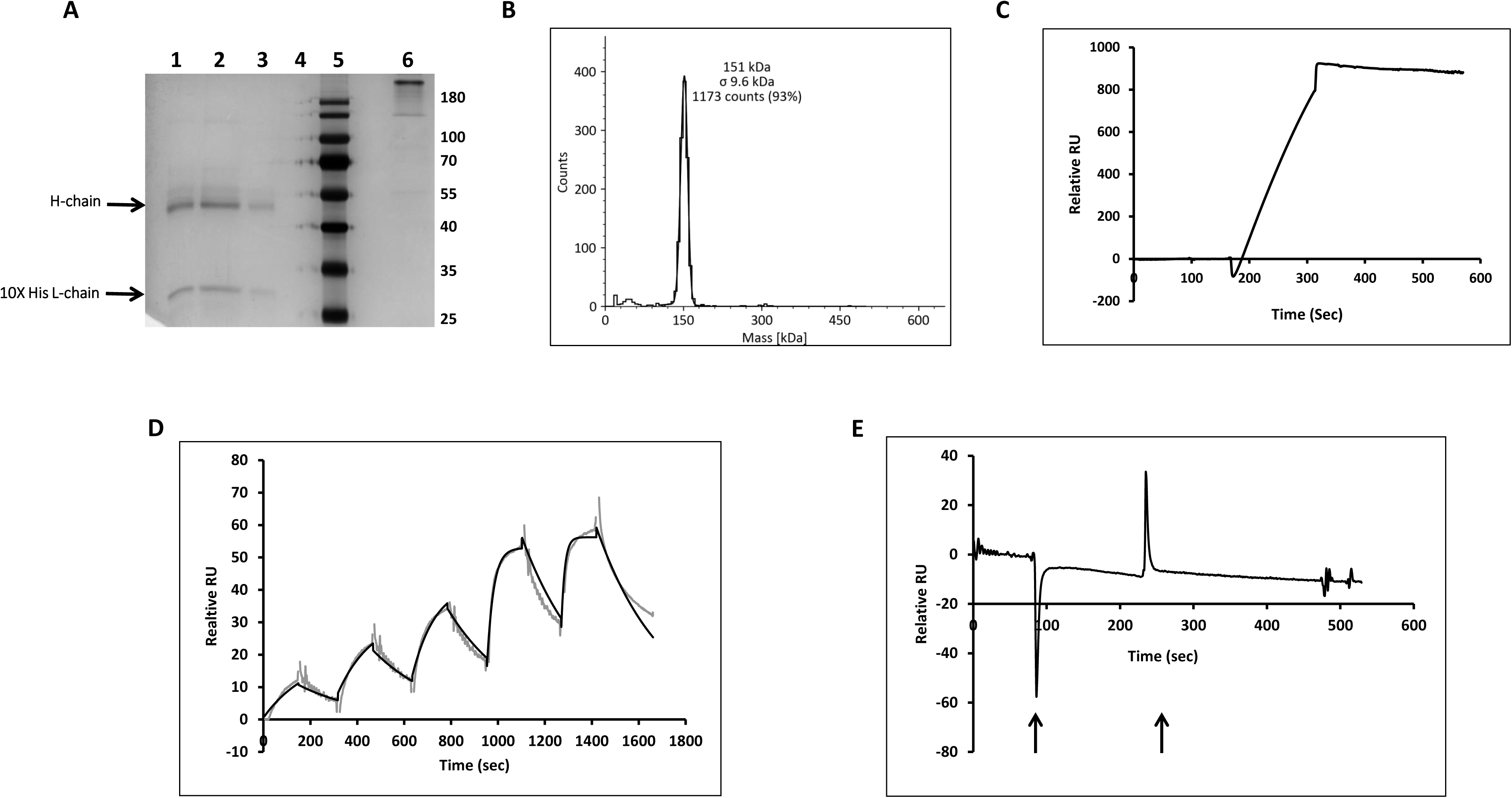
Measurement of binding affinity of His-10 chimeric 238A1 mAb using SPR on an NTA sensor chip: (A) Polyacrylamide gel shows migration of antibody heavy and light chains in different eluent fractions (50-, 100-, 200- and 300-mM imidazole) of IMAC purified His-10 chimeric 238A1 mAb under reducing conditions (*lanes 1-4*). Migration of the major eluent fraction of the mAb under non-reducing condition is shown in *lane 6*. (B) Histogram represents the molecular mass distribution of His-10 chimeric 238A1 mAb. (C) Sensorgram represents immobilization of His-10 mAb on NTA sensor chip in real-time, where purified antibody (0.1 nM) in HBS-P+ buffer was passed over the NTA chip. (D) A representative single cycle kinetic analysis of direct binding of monovalent analyte, Bio-pVIII peptide complexed with streptavidin, and His-10 chimeric 238A1 mAb immobilized on an NTA chip is shown. The experimental data are shown in *gray* and the calculated fit using a 1:1 binding model is shown in *black*. Affinity measurements were conducted using SPR in 3 independent binding reactions and the K_d_ value (mentioned in text) is represented as mean ± standard deviation. (E) Sensorgram showing interaction of biotin blocked streptavidin control analyte with immobilized His-10 238A1 mAb. Two spikes, shown by arrows, represent sudden refractive index changes at the start and end of sample injection.

The His-10 chimeric mAb showed a single major peak corresponding to 151 ± 9.6 kDa which suggests that chimerization or incorporation of His-10 at the C-terminus of the light chain did not affect the monomeric status of 238A1 mAb (Fig 9B). The determination of the antigen-antibody affinity constant is an important component of the characterization of a newly discovered antibody. SPR is a rapid and reliable method for measuring the strength of biomolecular interactions. The analysis typically requires the immobilization of IgG on the biosensor chip and flowing monovalent antigen over the chip surface. This approach is meant to avoid confounding effects on the binding data of avidity and rebinding by bivalent IgG. The immobilization of antibody mostly involves amine coupling or biotinylation of antibodies, which places them in a random orientation on the chip. To avoid these issues and provide new modes of antibody immobilization, alternative methods include Protein A-mediated binding or using a secondary antibody to immobilize the antibody of interest on the sensor chip. To further contribute to methods for stable antibody immobilization, we explored whether His-10 chimeric IgG molecules offer a simple alternative for oriented and reversible immobilization of antibodies on a NTA sensor chip. To this end, we immobilized purified His-10 238A1 mAb in the Flow Cell 2 of a NTA sensor chip, whereas His-10 17/9 chimeric mAb was immobilized in Flow Cell 1 as a non-specific reference following the manufacturers’ recommendations. The observed SPR signal derived from chip immobilization of His-10 chimeric mAb was found to be robust and stable even upon buffer flow after injection (Fig 9C). This shows that His-10 chimeric antibody can be stably immobilized on the NTA chip and can be used for affinity measurements in a 1:1 Langmuir binding mode.

The affinity constant for the His-10 238A1 mAb was determined for complexes of the Bio-pVIII peptide and streptavidin; the latter complexes were generated with a 20-fold molar excess of biotin to ensure monovalency of the antigen (i.e., only one or less biotin binding site per strepatvidin tetramer was occupied by a Bio-pVIII peptide). These streptavidin-peptide complexes were used as analyte to ensure both 1:1 binding and to obtain an appreciable signal-over-noise ratio. A biotin-blocked streptavidin analyte served as control. The passage of streptavidin-peptide complex, at different concentrations in a single cycle kinetics mode, generated clear resonance signals with distinct association and dissociation phases. The binding data were fit to a Langmuir 1:1 binding model and K_d_ value for the interaction under these conditions was 29.1 nM ± 2.8 nM for the His-10 238A1 antibody (Fig 9D). In contrast, flow of the biotin-blocked control analyte over the antibody immobilized sensor chip, even at highest streptavidin concentration (500 nM), did not generate a noticeable signal (Fig 9E). These results show that this cloning strategy resulted in the discovery of a rabbit-human chimeric mAb from a few single antigen-specific B cells.

## Discussion

The work described here proposes a facile pipeline for rapid screening and characterization of antibody Fv domains usually obtained from various antibody discovery workflows. We re-engineered a genetic system for generation of chimeric mAb expression cassettes that demonstrated reliable expression in mammalian culture. We also explored the possibility of using polyhistidine tags for purification and characterization of chimeric mAb forms of Fv domains and integrated it into our pipeline. The presence of a His-tag at the C-terminus of either the heavy or light chain did not influence the expression of chimeric mAbs in mammalian cells; however, for IMAC purification purposes, the optimal location for incorporation of His-10 into a mAb is at the C-terminus of the kappa light chain. In addition, our data show that the presence of His-6 at that location is not sufficient to facilitate affinity purification; the extended His-10 is required. It was also observed that presence of either His-6 or His-10 at the C-terminus of the gamma-1 heavy chain did not allow mAb purification. Some possible reasons for the inefficiency of these non-optimal his-tags may be their occlusion in the folded structure or their degradation. Our data do not provide evidence clarifying the specific mechanism behind the non-optimal performance of certain his-tags; that would require further study. Our observations also indicate that incorporation of the optimal His-10 to the C_κ_ chain does not strongly affect antigen binding affinity of the mAbs. Taken together, these observations suggest that the incorporation of a His-10 tag at the C-terminus C_κ_ chain provides the best way to facilitate one-step enrichment/purification of chimeric mAbs using IMAC.

Antibodies from humans and other mammals are composed of one of the five possible heavy chains (µ, δ, γ, ɑ or ε) and one of the two possible light chains (κ and λ). The constant region of both human light chains are similar in length (Cκ = 107 aa; C_λ_ = 105 aa) [63], overall structure and their interaction with the heavy chain. The range in the ratio of κ to λ usage in human serum antibodies varies from 0.85-1.86 [64]. On the other hand, rodents (20:1) and rabbits strongly favor kappa light chains while horses, cows, dogs and cats strongly favor lambda [65, 66]. The primary reason for selecting the human Cκ region for incorporating His-10 is that the human kappa locus, on chromosome 2, has only one constant region, whereas the human lambda locus has four possible functional constant regions [63]. The exclusivity of human Cκ region makes it a better starting point for developing a generic expression and purification system for human chimeric mAbs. Nonetheless, further studies are required to assess the potential of different human C_λ_ genes to support His-tag-based purification of bivalent human chimeric mAbs.

As a part of the validation for integration of our Fv domain cloning and purification system in antibody discovery pipeline, our initial effort led to the discovery of two novel, rabbit antibody genes specific for the filamentous phage major coat protein, pVIII. Due to the lack of well-characterized cell surface markers, our strategy of sorting antigen-specific rabbit B cells has relied mostly on positive antigen staining, Fc staining to ensure the presence of class switched IgG antibody, and decoy staining with biotin blocked streptavidin for removal of non-specific binders. However, the efficiency of selecting single, antigen-specific B cells may be improved further by negatively staining and discarding IgM^+^ B cells and CD4^+^ or CD8^+^ T cells as described by Starkie *et al.* [67]. The newly identified, natively paired rabbit V_H_ and V_L_ gene fragments were reformatted to full-length rabbit-human chimeric heavy and light chain genes. Anti-pVIII antibody, 238A1, uses the rabbit IGVH1S40 germline gene, which is in accordance with previous reports describing prevalent usage of this V_H_ gene segment in the rabbit antibody repertoire [68, 69], whereas the rabbit light chain variable germline gene used in this chimeric mAb is IGKV1S37.

We also analyzed the possible effect of chimerization and His-10 incorporation in our Fv cloning pipeline. The comparison of binding affinity values of mouse 8H11 and human chimeric 8H11 suggests that chimerization led to a roughly 40% reduction in affinity for 8H11. This could be due to some change in the conformation of critically important affinity determinants in the human chimeric version of 8H11mAb. It means that adoption of our approach for human chimerization may cause a certain decrease in native antibody affinity, depending on the degree of influence of the human constant regions on the paratope conformation of antibodies under study. The incorporation of His-10 also has certain shortcomings. Addition of His-10 to the carboxyl terminus of light chain further decreased antibody affinity by ∼20% in the two cases we studied. This reduction may be due to some interference of His-10 on light and heavy chain association.

It is likely that our genetic system of chimerization and His-10 incorporation may affect antigen-antibody affinities differently and cause a more drastic affinity reduction of less stable Fv domains among the subset under study in an antibody discovery workflow. Nevertheless, adoption of our cloning pipeline in antibody projects, involving discovery of non-human Fv domains, provides a rapid way to screen antibodies and obtain a reasonable estimate of their affinities. In addition, given the fact that incorporation of His-10 showed only a minor effect, our cloning system is less likely to affect native Fv domain affinities in projects involving antibody discovery from human origin, where relatively fewer changes in constant region is involved. Some of the commonly used strategies for discovering mAbs of human origin include immunizing transgenic mice encoding human immunoglobulin germline gene loci or using human antibody phage display libraries [70–72].

Incorporation of His-10 into a IgG1 mAb, however, also offers several advantages. First, His-10 recombinant mAbs can be generically purified using economical, robust, reusable, and rechargeable resins which may result in significant cost benefits in scale up operations. The recombinant mAbs can be purified or enriched directly from serum-supplemented, mammalian cell culture media in a single step. Second, the elution of these recombinant mAbs from the column does not require exposure to low pH buffers minimizing the possibility of activity loss or aggregation. Furthermore, the efficient resin sanitization after the column use for next round of charging can be performed using relatively harsher chemicals without significantly compromising the column binding capacity of IMAC resins. Third, His-10 recombinant bivalent antibodies can be directly immobilized by passing cell culture media over an NTA sensor chip for affinity measurements without the need to purify the mAbs. The affinity measurements of these mAbs can be reliably performed on Ni-NTA biosensors without much interference due to rebinding or avidity effects following a simple 1:1 binding mode. Finally, the His-10 in chimeric mAbs can be exploited for detection of biomolecules or interacting partners in various immunological analysis.

Overall, our genetic system simplifies reconstruction of Fv fragments derived from single B cells into His-10 human chimeric bivalent mAbs. The expression of reformatted His-10 antibodies facilitates generic purification of chimeric mAbs. Moreover, the mAb format provides a rapid and reasonable estimate of binding affinities of novel bivalent chimeric mAbs. The advantages associated with expression of His-10 recombinant mAbs offer great potential for its incorporation in novel or established antibody discovery pipelines.

## Author contributions

JKS supervised the study. AS, NG and JKS designed experiments. AS performed the experiments with assistance from DP. AS, NG, JMP and JKS analyzed the data. TJW provided critical suggestions for the manuscript. AS and JKS wrote the manuscript which all authors read, edited, and revised.

## Acknowledgements

We thank Tim Heslip for his help with FACS instrumentation and members of the laboratories of Drs. Jonathan Choy, Z. Brumme, and M. Brockman for their suggestions. We are grateful to the Office of Animal Care Services at Simon Fraser University for their help with rabbit care, immunizations, and sample collection. We also acknowledge the contributions of Dr. Nicholas Schnicker at the Protein and Crystallography facility in the University of Iowa with BLI instrumentation.

## Availability of antibody expression cassettes

The functional antibody expression cassettes used in this study is available for use to the research community upon request. We, however, need to seek permission from the Duke university before distribution.

## Abbreviations

Bio-HA: Biotinylated hemagglutinin peptide
Bio-pVIII: Biotinylated filamentous phage pVIII peptide
BLI: Biolayer interferometry
BSA: Bovine serum albumin
DMSO: Dimethyl sulfoxide
ECD: Extracellular domain
ELISA: Enzyme-linked immunosorbent assay
FACS: Fluorescence-activated cell sorting
FBS: Fetal bovine serum
HA: Hemagglutinin peptide
HRP: Horseradish peroxidase
HS-TBS: High-strength tris buffered saline
IMAC: Immobilized metal-chelate affinity chromatography
mAbs: Monoclonal antibodies
Ni: Nickel
NTA: Nitrilotriacetic acid
NZW: New Zealand White
pVIII: Major coat protein of filamentous bacteriophage
PBS: Phosphate buffered saline
P_CMV_: Cytomegalovirus promoter
SP: Signal peptide
SPR: Surface plasmon resonance
TBS: Tris buffered saline

## References

1 Pedrioli, A & Oxenius, A (2021) Single B cell technologies for monoclonal antibody discovery. Trends Immunol 42: 1143–1158.

2 Lu, RM, Hwang, YC, Liu, IJ, Lee, CC, Tsai, HZ, Li, HJ & Wu, HC (2020) Development of therapeutic antibodies for the treatment of diseases. J Biomed Sci 27: 1.

3 Kohler, G & Milstein, C (1975) Continuous cultures of fused cells secreting antibody of predefined specificity. Nature 256: 495–497.

4 Smith, SA & Crowe, JE, Jr. (2015) Use of Human Hybridoma Technology To Isolate Human Monoclonal Antibodies. Microbiol Spectr 3: AID-0027-2014.

5 Knezevic, I, Kang, HN & Thorpe, R (2015) Immunogenicity assessment of monoclonal antibody products: A simulated case study correlating antibody induction with clinical outcomes. Biologicals 43: 307–317.

6 Wardemann, H, Yurasov, S, Schaefer, A, Young, JW, Meffre, E & Nussenzweig, MC (2003) Predominant autoantibody production by early human B cell precursors. Science 301: 1374–1377.

7 Tiller, T, Meffre, E, Yurasov, S, Tsuiji, M, Nussenzweig, MC & Wardemann, H (2008) Efficient generation of monoclonal antibodies from single human B cells by single cell RT-PCR and expression vector cloning. J Immunol Methods 329: 112–124.

8 Tiller, T, Busse, CE & Wardemann, H (2009) Cloning and expression of murine Ig genes from single B cells. J Immunol Methods 350: 183–193.

9 Volkheimer, AD, Weinberg, JB, Beasley, BE, Whitesides, JF, Gockerman, JP, Moore, JO, Kelsoe, G, Goodman, BK & Levesque, MC (2007) Progressive immunoglobulin gene mutations in chronic lymphocytic leukemia: evidence for antigen-driven intraclonal diversification. Blood 109: 1559–1567.

10 Smith, K, Garman, L, Wrammert, J, Zheng, NY, Capra, JD, Ahmed, R & Wilson, PC (2009) Rapid generation of fully human monoclonal antibodies specific to a vaccinating antigen. Nat Protoc 4: 372–384.

11 Wrammert, J, Smith, K, Miller, J, Langley, WA, Kokko, K, Larsen, C, Zheng, NY, Mays, I, Garman, L, Helms, C, James, J, Air, GM, Capra, JD, Ahmed, R & Wilson, PC (2008) Rapid cloning of high-affinity human monoclonal antibodies against influenza virus. Nature 453: 667–671.

12 Wang, B, Kluwe, CA, Lungu, OI, DeKosky, BJ, Kerr, SA, Johnson, EL, Tanno, H, Lee, CH, Jung, J, Rezigh, AB, Carroll, SM, Reyes, AN, Bentz, JR, Villanueva, I, Altman, AL, Davey, RA, Ellington, AD & Georgiou, G (2015) Facile Discovery of a Diverse Panel of Anti-Ebola Virus Antibodies by Immune Repertoire Mining. Sci Rep 5: 13926.

13 DeKosky, BJ, Kojima, T, Rodin, A, Charab, W, Ippolito, GC, Ellington, AD & Georgiou, G (2015) In-depth determination and analysis of the human paired heavy- and light-chain antibody repertoire. Nat Med 21: 86–91.

14 DeKosky, BJ, Ippolito, GC, Deschner, RP, Lavinder, JJ, Wine, Y, Rawlings, BM, Varadarajan, N, Giesecke, C, Dorner, T, Andrews, SF, Wilson, PC, Hunicke-Smith, SP, Willson, CG, Ellington, AD & Georgiou, G (2013) High-throughput sequencing of the paired human immunoglobulin heavy and light chain repertoire. Nat Biotechnol 31: 166–169.

15 Rowinsky, EK, Youssoufian, H, Tonra, JR, Solomon, P, Burtrum, D & Ludwig, DL (2007) IMC-A12, a human IgG1 monoclonal antibody to the insulin-like growth factor I receptor. Clin Cancer Res 13: 5549s–5555s.

16 Bedinger, D, Lao, L, Khan, S, Lee, S, Takeuchi, T & Mirza, AM (2016) Development and characterization of human monoclonal antibodies that neutralize multiple TGFbeta isoforms. MAbs 8: 389–404.

17 Pukac, L, Kanakaraj, P, Humphreys, R, Alderson, R, Bloom, M, Sung, C, Riccobene, T, Johnson, R, Fiscella, M, Mahoney, A, Carrell, J, Boyd, E, Yao, XT, Zhang, L, Zhong, L, von Kerczek, A, Shepard, L, Vaughan, T, Edwards, B, Dobson, C, Salcedo, T & Albert, V (2005) HGS-ETR1, a fully human TRAIL-receptor 1 monoclonal antibody, induces cell death in multiple tumour types in vitro and in vivo. Br J Cancer 92: 1430–1441.

18 Burmester, GR, Feist, E, Sleeman, MA, Wang, B, White, B & Magrini, F (2011) Mavrilimumab, a human monoclonal antibody targeting GM-CSF receptor-alpha, in subjects with rheumatoid arthritis: a randomised, double-blind, placebo-controlled, phase I, first-in-human study. Ann Rheum Dis 70: 1542–1549.

19 McMahon, C, Baier, AS, Pascolutti, R, Wegrecki, M, Zheng, S, Ong, JX, Erlandson, SC, Hilger, D, Rasmussen, SGF, Ring, AM, Manglik, A & Kruse, AC (2018) Yeast surface display platform for rapid discovery of conformationally selective nanobodies. Nat Struct Mol Biol 25: 289–296.

20 Fisher, TS, Kamperschroer, C, Oliphant, T, Love, VA, Lira, PD, Doyonnas, R, Bergqvist, S, Baxi, SM, Rohner, A, Shen, AC, Huang, C, Sokolowski, SA & Sharp, LL (2012) Targeting of 4-1BB by monoclonal antibody PF-05082566 enhances T-cell function and promotes anti-tumor activity. Cancer Immunol Immunother 61: 1721–1733.

21 Bohrmann, B, Baumann, K, Benz, J, Gerber, F, Huber, W, Knoflach, F, Messer, J, Oroszlan, K, Rauchenberger, R, Richter, WF, Rothe, C, Urban, M, Bardroff, M, Winter, M, Nordstedt, C & Loetscher, H (2012) Gantenerumab: a novel human anti-Abeta antibody demonstrates sustained cerebral amyloid-beta binding and elicits cell-mediated removal of human amyloid-beta. J Alzheimers Dis 28: 49–69.

22 Villa, A, Trachsel, E, Kaspar, M, Schliemann, C, Sommavilla, R, Rybak, JN, Rosli, C, Borsi, L & Neri, D (2008) A high-affinity human monoclonal antibody specific to the alternatively spliced EDA domain of fibronectin efficiently targets tumor neo-vasculature in vivo. Int J Cancer 122: 2405–2413.

23 Lach-Trifilieff, E, Minetti, GC, Sheppard, K, Ibebunjo, C, Feige, JN, Hartmann, S, Brachat, S, Rivet, H, Koelbing, C, Morvan, F, Hatakeyama, S & Glass, DJ (2014) An antibody blocking activin type II receptors induces strong skeletal muscle hypertrophy and protects from atrophy. Mol Cell Biol 34: 606–618.

24 Chowdhury, PS, Viner, JL, Beers, R & Pastan, I (1998) Isolation of a high-affinity stable single-chain Fv specific for mesothelin from DNA-immunized mice by phage display and construction of a recombinant immunotoxin with anti-tumor activity. Proc Natl Acad Sci U S A 95: 669–674.

25 Hassan, R, Ebel, W, Routhier, EL, Patel, R, Kline, JB, Zhang, J, Chao, Q, Jacob, S, Turchin, H, Gibbs, L, Phillips, MD, Mudali, S, Iacobuzio-Donahue, C, Jaffee, EM, Moreno, M, Pastan, I, Sass, PM, Nicolaides, NC & Grasso, L (2007) Preclinical evaluation of MORAb-009, a chimeric antibody targeting tumor-associated mesothelin. Cancer Immun 7: 20.

26 Bakker, AB, Marissen, WE, Kramer, RA, Rice, AB, Weldon, WC, Niezgoda, M, Hanlon, CA, Thijsse, S, Backus, HH, de Kruif, J, Dietzschold, B, Rupprecht, CE & Goudsmit, J (2005) Novel human monoclonal antibody combination effectively neutralizing natural rabies virus variants and individual in vitro escape mutants. J Virol 79: 9062–9068.

27 Kramer, RA, Marissen, WE, Goudsmit, J, Visser, TJ, Clijsters-Van der Horst, M, Bakker, AQ, de Jong, M, Jongeneelen, M, Thijsse, S, Backus, HH, Rice, AB, Weldon, WC, Rupprecht, CE, Dietzschold, B, Bakker, AB & de Kruif, J (2005) The human antibody repertoire specific for rabies virus glycoprotein as selected from immune libraries. Eur J Immunol 35: 2131–2145.

28 Liao, HX, Levesque, MC, Nagel, A, Dixon, A, Zhang, R, Walter, E, Parks, R, Whitesides, J, Marshall, DJ, Hwang, KK, Yang, Y, Chen, X, Gao, F, Munshaw, S, Kepler, TB, Denny, T, Moody, MA & Haynes, BF (2009) High-throughput isolation of immunoglobulin genes from single human B cells and expression as monoclonal antibodies. J Virol Methods 158: 171–179.

29 Shukla, AA, Gupta, P & Han, X (2007) Protein aggregation kinetics during Protein A chromatography. Case study for an Fc fusion protein. J Chromatogr A 1171: 22–28.

30 Xu, X, Didio, DM, Leister, KJ & Ghose, S (2010) Disaggregation of high-molecular weight species during downstream processing to recover functional monomer. Biotechnol Prog 26: 717–726.

31 Fuglistaller, P (1989) Comparison of immunoglobulin binding capacities and ligand leakage using eight different protein A affinity chromatography matrices. J Immunol Methods 124: 171–177.

32 Ramos-de-la-Pena, AM, Gonzalez-Valdez, J & Aguilar, O (2019) Protein A chromatography: Challenges and progress in the purification of monoclonal antibodies. J Sep Sci 42: 1816–1827.

33 Yamada, T, Yamamoto, K, Ishihara, T & Ohta, S (2017) Purification of monoclonal antibodies entirely in flow-through mode. J Chromatogr B Analyt Technol Biomed Life Sci 1061-1062: 110–116.

34 Naik, AD, Menegatti, S, Gurgel, PV & Carbonell, RG (2011) Performance of hexamer peptide ligands for affinity purification of immunoglobulin G from commercial cell culture media. J Chromatogr A 1218: 1691–1700.

35 Verdoliva, A, Pannone, F, Rossi, M, Catello, S & Manfredi, V (2002) Affinity purification of polyclonal antibodies using a new all-D synthetic peptide ligand: comparison with protein A and protein G. J Immunol Methods 271: 77–88.

36 Mao, LN, Rogers, JK, Westoby, M, Conley, L & Pieracci, J (2010) Downstream antibody purification using aqueous two-phase extraction. Biotechnol Prog 26: 1662–1670.

37 Wu, Q, Lin, DQ, Zhang, QL, Gao, D & Yao, SJ (2014) Evaluation of a PEG/hydroxypropyl starch aqueous two-phase system for the separation of monoclonal antibodies from cell culture supernatant. J Sep Sci 37: 447–453.

38 Kang, YK, Hamzik, J, Felo, M, Qi, B, Lee, J, Ng, S, Liebisch, G, Shanehsaz, B, Singh, N, Persaud, K, Ludwig, DL & Balderes, P (2013) Development of a novel and efficient cell culture flocculation process using a stimulus responsive polymer to streamline antibody purification processes. Biotechnol Bioeng 110: 2928–2937.

39 Singh, N, Pizzelli, K, Romero, JK, Chrostowski, J, Evangelist, G, Hamzik, J, Soice, N & Cheng, KS (2013) Clarification of recombinant proteins from high cell density mammalian cell culture systems using new improved depth filters. Biotechnol Bioeng 110: 1964–1972.

40 Capito, F, Skudas, R, Stanislawski, B & Kolmar, H (2013) Customization of copolymers to optimize selectivity and yield in polymer-driven antibody purification processes. Biotechnol Prog 29: 1484–1493.

41 Kimple, ME, Brill, AL & Pasker, RL (2013) Overview of affinity tags for protein purification. Curr Protoc Protein Sci 73: 9.9.1–9.9.23.

42 Crowe, J, Dobeli, H, Gentz, R, Hochuli, E, Stuber, D & Henco, K (1994) 6xHis-Ni-NTA chromatography as a superior technique in recombinant protein expression/purification. Methods Mol Biol 31: 371–387.

43 Hale, JE & Beidler, DE (1994) Purification of humanized murine and murine monoclonal antibodies using immobilized metal-affinity chromatography. Anal Biochem 222: 29–33.

44 Boden, V, Winzerling, JJ, Vijayalakshmi, M & Porath, J (1995) Rapid one-step purification of goat immunoglobulins by immobilized metal ion affinity chromatography. J Immunol Methods 181: 225–232.

45 Porath, J & Olin, B (1983) Immobilized metal ion affinity adsorption and immobilized metal ion affinity chromatography of biomaterials. Serum protein affinities for gel-immobilized iron and nickel ions. Biochemistry 22: 1621–1630.

46 Pham, PL, Perret, S, Doan, HC, Cass, B, St-Laurent, G, Kamen, A & Durocher, Y (2003) Large-scale transient transfection of serum-free suspension-growing HEK293 EBNA1 cells: peptone additives improve cell growth and transfection efficiency. Biotechnol Bioeng 84: 332–342.

47 Ill, CR, Brehm, T, Lydersen, BK, Hernandez, R & Burnett, KG (1988) Species specificity of iron delivery in hybridomas. In Vitro Cell Dev Biol 24: 413–419.

48 Ill, CR, Keivens, VM, Hale, JE, Nakamura, KK, Jue, RA, Cheng, S, Melcher, ED, Drake, B & Smith, MC (1993) A COOH-terminal peptide confers regiospecific orientation and facilitates atomic force microscopy of an IgG1. Biophys J 64: 919–924.

49 Li, W, Fan, Z, Lin, Y & Wang, TY (2021) Serum-Free Medium for Recombinant Protein Expression in Chinese Hamster Ovary Cells. Front Bioeng Biotechnol 9: 646363.

50 Cotten, M, Stegmueller, K, Eickhoff, J, Hanke, M, Herzberger, K, Herget, T, Choidas, A, Daub, H & Godl, K (2003) Exploiting features of adenovirus replication to support mammalian kinase production. Nucleic Acids Res 31: e128.

51 Sardy, M, Odenthal, U, Karpati, S, Paulsson, M & Smyth, N (1999) Recombinant human tissue transglutaminase ELISA for the diagnosis of gluten-sensitive enteropathy. Clin Chem 45: 2142–2149.

52 Khawli, LA, Biela, BH, Hu, P & Epstein, AL (2002) Stable, genetically engineered F(ab’)(2) fragments of chimeric TNT-3 expressed in mammalian cells. Hybrid Hybridomics 21: 11–18.

53 Voss, S & Skerra, A (1997) Mutagenesis of a flexible loop in streptavidin leads to higher affinity for the Strep-tag II peptide and improved performance in recombinant protein purification. Protein Eng 10: 975–982.

54 Maertens, B, Spriestersbach, A, Kubicek, J & Schafer, F (2015) Strep-Tagged Protein Purification. Methods Enzymol 559: 53–69.

55 Findlay, JW & Dillard, RF (2007) Appropriate calibration curve fitting in ligand binding assays. AAPS J 9: E260–267.

56 Burstein, Y & Schechter, I (1978) Primary structures of N-terminal extra peptide segments linked to the variable and constant regions of immunoglobulin light chain precursors: implications on the organization and controlled expression of immunoglobulin genes. Biochemistry 17: 2392–2400.

57 Park, JM, Yang, X, Park, JJ, Press, OW & Press, MF (1999) Assessment of novel anti-p185HER-2 monoclonal antibodies for internalization-dependent therapies. Hybridoma 18: 487–495.

58 Wan, L, Zhu, S, Zhu, J, Yang, H, Li, S, Li, Y, Cheng, J & Lu, X (2013) Production and characterization of a CD25-specific scFv-Fc antibody secreted from Pichia pastoris. Appl Microbiol Biotechnol 97: 3855–3863.

59 Powers, DB, Amersdorfer, P, Poul, M, Nielsen, UB, Shalaby, MR, Adams, GP, Weiner, LM & Marks, JD (2001) Expression of single-chain Fv-Fc fusions in Pichia pastoris. J Immunol Methods 251: 123–135.

60 Schulze-Gahmen, U, Rini, JM, Arevalo, J, Stura, EA, Kenten, JH & Wilson, IA (1988) Preliminary crystallographic data, primary sequence, and binding data for an anti-peptide Fab and its complex with a synthetic peptide from influenza virus hemagglutinin. J Biol Chem 263: 17100–17105.

61 Niman, HL, Houghten, RA, Walker, LE, Reisfeld, RA, Wilson, IA, Hogle, JM & Lerner, RA (1983) Generation of protein-reactive antibodies by short peptides is an event of high frequency: implications for the structural basis of immune recognition. Proc Natl Acad Sci U S A 80: 4949–4953.

62 Giudicelli, V, Chaume, D & Lefranc, MP (2005) IMGT/GENE-DB: a comprehensive database for human and mouse immunoglobulin and T cell receptor genes. Nucleic Acids Res 33: D256–261.

63 Solomon, A & Weiss, DT (1995) Structural and functional properties of human lambda-light-chain variable-region subgroups. Clin Diagn Lab Immunol 2: 387–394.

64 Barandun, S, Morell, A, Skvaril, F & Oberdorfer, A (1976) Deficiency of kappa- or lambda-type immunoglobulins. Blood 47: 79–89.

65 Arun, SS, Breuer, W & Hermanns, W (1996) Immunohistochemical examination of light-chain expression (lambda/kappa ratio) in canine, feline, equine, bovine and porcine plasma cells. Zentralbl Veterinarmed A 43: 573–576.

66 Woloschak, GE & Krco, CJ (1987) Regulation of kappa/lambda immunoglobulin light chain expression in normal murine lymphocytes. Mol Immunol 24: 751–757.

67 Starkie, DO, Compson, JE, Rapecki, S & Lightwood, DJ (2016) Generation of Recombinant Monoclonal Antibodies from Immunised Mice and Rabbits via Flow Cytometry and Sorting of Antigen-Specific IgG+ Memory B Cells. PLoS One 11: e0152282.

68 Knight, KL (1992) Restricted VH gene usage and generation of antibody diversity in rabbit. Annu Rev Immunol 10: 593–616.

69 Kodangattil, S, Huard, C, Ross, C, Li, J, Gao, H, Mascioni, A, Hodawadekar, S, Naik, S, Min-debartolo, J, Visintin, A & Almagro, JC (2014) The functional repertoire of rabbit antibodies and antibody discovery via next-generation sequencing. MAbs 6: 628–636.

70 Bruggemann, M, Caskey, HM, Teale, C, Waldmann, H, Williams, GT, Surani, MA & Neuberger, MS (1989) A repertoire of monoclonal antibodies with human heavy chains from transgenic mice. Proc Natl Acad Sci U S A 86: 6709–6713.

71 Bruggemann, M, Spicer, C, Buluwela, L, Rosewell, I, Barton, S, Surani, MA & Rabbitts, TH (1991) Human antibody production in transgenic mice: expression from 100 kb of the human IgH locus. Eur J Immunol 21: 1323–1326.

72 McCafferty, J, Griffiths, AD, Winter, G & Chiswell, DJ (1990) Phage antibodies: filamentous phage displaying antibody variable domains. Nature 348: 552–554.

