## Supplementary Fig 1 for "A pipeline for facile cloning of antibody Fv domains and their expression, purification, and characterization as recombinant His-tagged IgGs"

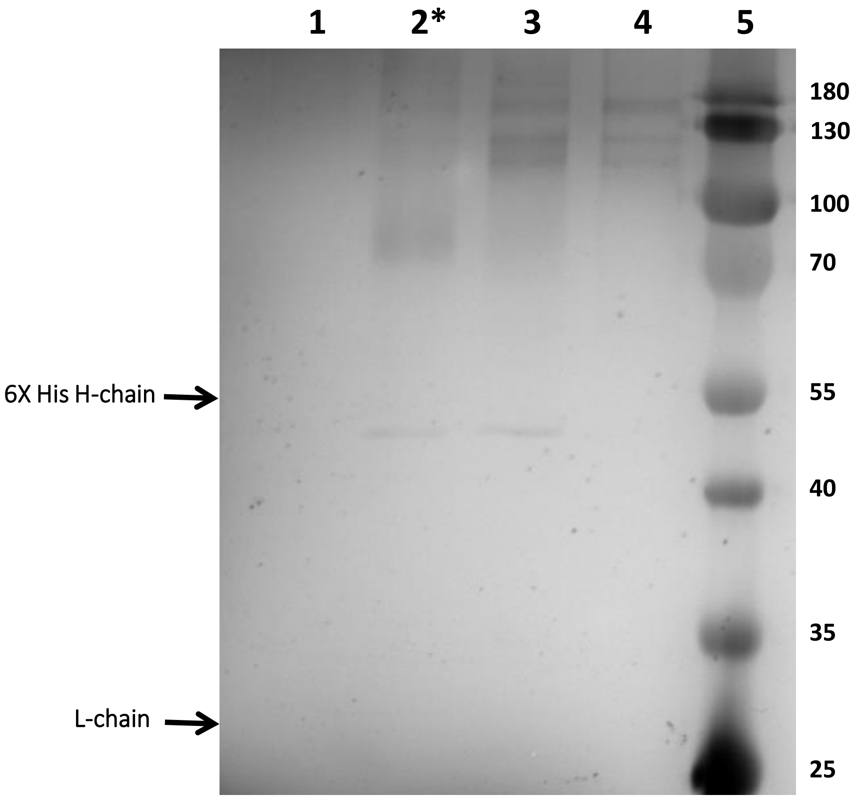


Sinha *et al.* Supplementary figure 1

**Supplementary figure 1 legend**

**SDS-PAGE gel exhibiting eluent fractions of Ni-NTA column obtained from culture supernatant expressing chimeric 8H11 containing His-6 heavy chain:** Silver stained 12% SDS-PAGE gel showing the different eluent fractions (50-, 100-, 200- and 300- mM) from Ni-NTA column derived from cell culture media expressing His-6 IgG1 heavy chain containing chimeric 8H11 mAb (*lanes 1-5*). The numbers indicated adjacent to the gel represent the molecular weight of polypeptides (in kDa) in the protein ladder. Specific heavy chain and light chain polypeptides migrate with an apparent molecular weight of ~55 kDa and ~27 kDa (expected positions of both heavy and light chains are marked with arrows). The fraction where His-10 chimeric mAbs are usually eluted is represented by an *asterisk*.
